# Mapping Cognitive Brain Functions at Scale

**DOI:** 10.1101/2020.05.14.097014

**Authors:** Pragathi Priyadharsini Balasubramani, Alejandro Ojeda, Gillian Grennan, Vojislav Maric, Hortense Le, Fahad Alim, Mariam Zafar-Khan, Juan Diaz-Delgado, Sarita Silveira, Dhakshin Ramanathan, Jyoti Mishra

## Abstract

A fundamental set of cognitive abilities enable humans to efficiently process goal-relevant information, suppress irrelevant distractions, maintain information in working memory, and act flexibly in different behavioral contexts. Yet, studies of human cognition and their underlying neural mechanisms usually evaluate these cognitive constructs in silos, instead of comprehensively in-tandem within the same individual. Here, we developed a scalable, mobile platform, “*BrainE*” (short for Brain Engagement), to rapidly assay several essential aspects of cognition simultaneous with wireless electroencephalography (EEG) recordings. Using *BrainE*, we rapidly assessed five aspects of cognition including (1) selective attention, (2) response inhibition, (3) working memory, (4) flanker interference and (5) emotion interference processing, in 102 healthy young adults. We evaluated stimulus encoding in all tasks using the EEG neural recordings, and isolated the cortical sources of the spectrotemporal EEG dynamics. Additionally, we used *BrainE* in a two-visit study in 24 young adults to investigate the reliability of the neuro-cognitive data as well as its plasticity to transcranial magnetic stimulation (TMS). We found that stimulus encoding on multiple cognitive tasks could be rapidly assessed, identifying common as well as distinct task processes in both sensory and cognitive control brain regions. Event related synchronization (ERS) in the theta (3-7 Hz) and alpha (8-12 Hz) frequencies as well as event related desynchronization (ERD) in the beta frequencies (13-30 Hz) were distinctly observed in each task. The observed ERS/ERD effects were overall anticorrelated. The two-visit study confirmed high test-retest reliability for both cognitive and neural data, and neural responses showed specific TMS protocol driven modulation. We also show that the global cognitive neural responses are sensitive to mental health symptom self-reports. This first study with the *BrainE* platform showcases its utility in studying neuro-cognitive dynamics in a rapid and scalable fashion.

**Highlights:**

- Rapid and scalable EEG recordings reveal common and distinct cortical activations across five core cognitive tasks.
- Data acquired across visits one-week-apart show high test-retest reliability for both cognitive and neural measurements.
- Evoked neural responses during emotion interference processing demonstrate specific short-term plasticity driven by type of neurostimulation.
- Cognitively evoked neural responses are sensitive to variations in mental health symptoms.

## Introduction

Healthy brains are wired to effectively and efficiently process information. These complex systems simultaneously ensure stability as well as flexibility, and reflect an essential capacity to adapt to constantly changing environmental and motivational contexts. This dynamic ability of human brains requiring multiple interacting mental operations is referred to as cognitive control (Badre, 2011; Lenartowicz et al., 2010; Luna et al., 2015). Cognitive control operations fundamentally include abilities for stimulus encoding as well as online maintenance of goal-relevant information (Gazzaley and Nobre, 2012), suppression of competing goal-irrelevant distractions and behaviors (Mishra *et al*., 2013), and a continuous evaluation of the accuracy of selected actions based on feedback (Posner and Rothbart, 2009; van Noordt and Segalowitz, 2012). Much research to-date has focused on studying these individual component processes of cognitive control in isolation in select human population cohort studies. Yet, studies rarely evaluate these multiple essential cognitive operations within the same individual, particularly investigating their common and distinct underlying neural features. Thus, there is a gap in the comprehensive understanding of the neural circuit dynamics that underlie diverse cognitive states within the same individual. This lack of knowledge has translational implications. Multiple aspects of cognition are significantly altered in a range of neuropsychiatric disorders (Millan *et al*., 2012), but the degree to which these abnormalities are specific to a particular cognitive/neural circuit; or occur across many cognitive operations and states remains unknown.

Here, we developed a scalable, mobile platform, *BrainE* (short for Brain Engagement), which aims at assessing cognitive control within and across humans, rapidly evaluating several integral cognitive processes simultaneously with electroencephalography (EEG) based neural recordings. In *BrainE*, we adopt standard cognitive assessments of attention, response inhibition, working memory, and distractor suppression in both non-emotional and emotional contexts, that are designed to be engaging and equally interpretable for individuals from diverse cultural backgrounds and across the lifespan. With the objective to make cognitive brain mapping scalable and accessible, we integrated non-invasive, mobile and semi-dry electrode EEG within *BrainE* for simultaneously acquiring cognitive behavioral data and neural signals. In this first *BrainE* study, we conduct cognitive brain mapping in healthy adult human subjects, investigating neural processes underlying stimulus encoding in multiple cognitive contexts. We also derive the cortical sources of the observed spectrotemporal neural dynamics. Additionally, in a second study, we present data from a two-visit experiment to assess the reliability of *BrainE* recordings, as well as the sensitivity of the cognitive neural markers to neuromodulation using transcranial magnetic stimulation (TMS).

## Methods

### Experimental Design

#### Mental Health Ratings

All participants completed subjective mental health self-reports using standard instruments: inattention and hyperactivity ratings were obtained on the ADHD Rating Scale (New York University and Massachusetts General Hospital. Adult ADHD-RS-IV* with Adult Prompts. 2003; : 9– 10), anxiety was measured using the Generalized Anxiety Disorder 7-item scale GAD-7 (Spitzer *et al*., 2006)), and depression was reported on the 9-item Patient Health Questionnaire (PHQ-9 (Kroenke, Spitzer and Williams, 2001). We also obtained demographic variables by self-report including, age, gender, race and ethnicity, socio-economic status measured on the Family Affluence Scale (Boudreau and Poulin, 2008), and any current/past history of clinical diagnoses and medications.

#### *BrainE* Neuro-Cognitive Assessments

Assessments were developed and deployed by NEAT Labs (Misra et al., 2018) on the Unity game engine. The Lab Streaming Layer (LSL, Kothe et al., 2019) protocol was used to time-stamp each stimulus/response event in each cognitive task. Study participants engaged with *BrainE* assessments on a Windows-10 laptop sitting at a comfortable viewing distance. Participants underwent the following cognitive assessment modules that were completed within a 35 min session. **Figure 1** shows the stimulus sequence in each task.

**Figure 1.**
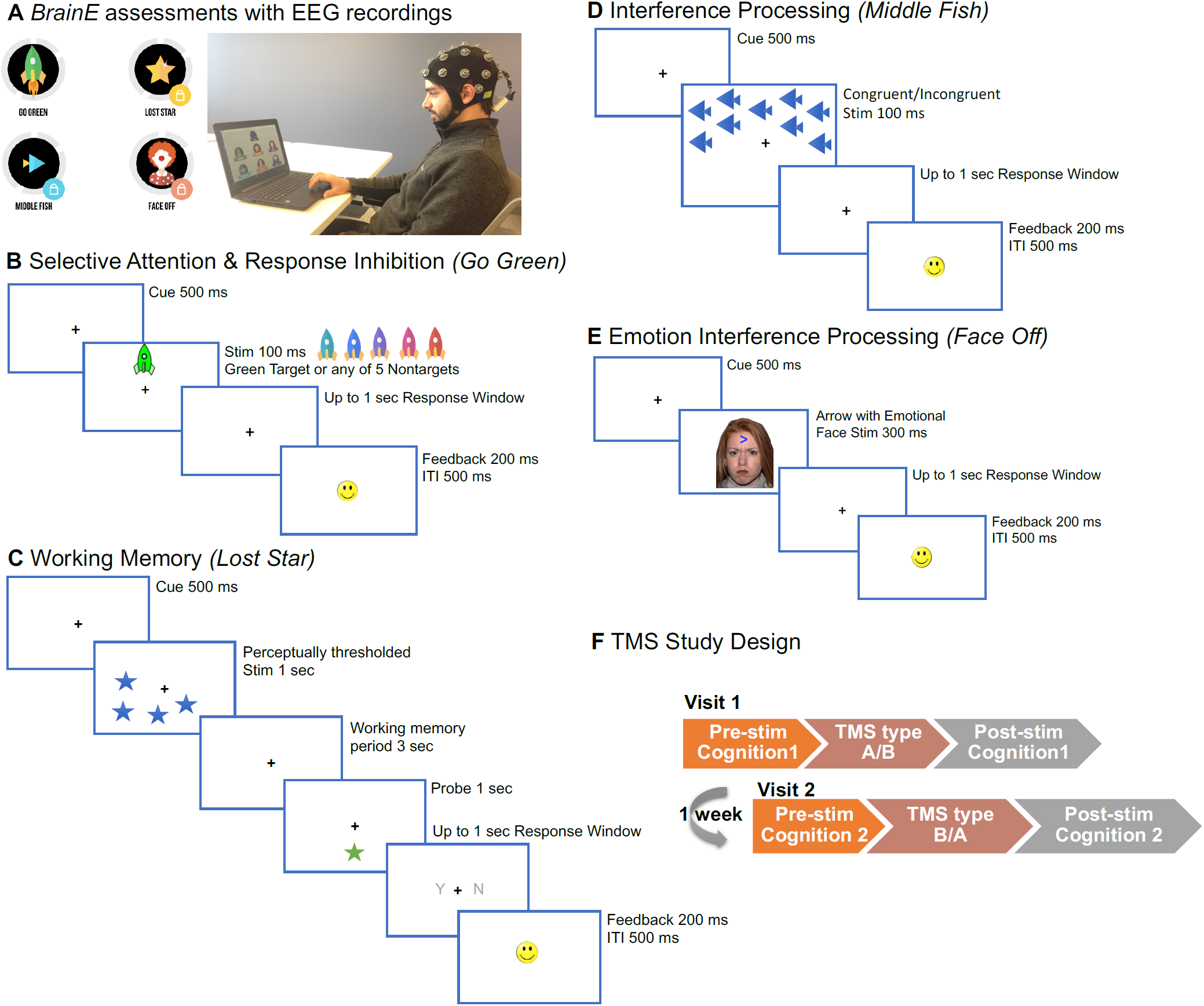
Cognitive studies delivered on the *BrainE* platform. (A) *BrainE* assessment dashboard with the wireless EEG recording setup. (B) The selective attention and response inhibition task differ only in the frequency of targets; sparse 33% targets appear in the Selective Attention block and frequent 67% targets appear in the Response Inhibition block. (C) Working memory task with perceptually thresholded stimuli. (D) Flanker interference processing task; flanking fish may either face the same direction as the middle fish on congruent trials, or the opposite direction on incongruent trials. (E) Emotion interference task presents neutral, happy, sad or angry faces superimposed on the arrow. (F) The TMS study involved two visits with two types of TMS stimulation A (cTBS) or B (iTBS) delivered in each week counterbalanced across subjects, and with immediate pre- and post-neurocognitive assessments.

##### 1. Selective Attention & Response Inhibition

Participants accessed a game named *Go Green* modeled after the standard test of variables of attention (Greenberg and Waldman, 1993). In this simple two-block task, colored rockets were presented either in the upper/lower central visual field. Participants were instructed to respond to green colored rocket targets and ignore, i.e. withhold their response to distracting rockets of five other isoluminant colors (shades of cyan, blue, purple, pink, orange). The task sequence consisted of a central fixation ‘+’ cue for 500 msec followed by a target/non-target stimulus of 100 msec duration, and up to a 1 sec duration blank response window. When the participant made a response choice, or at the end of 1 sec in case of no response, a happy or sad face emoticon was presented for 200 msec to signal response accuracy, followed by a 500 msec inter-trial interval (ITI). To reinforce positive feedback for fast and accurate responding, within 100-400 msec, two happy face emoticons were simultaneously presented during the feedback period (Wodka *et al*., 2007). Both task blocks had 90 trials lasting 5 min each, with target/non-target trials shuffled in each block. A brief practice period of 4 trials preceded the main task blocks. Summary total block accuracy was provided to participants at the end of each block as a series of happy face emoticons (up to 10 emoticons) in this and in all assessments described below.

In the first task block, green rocket targets were sparse (33% of trials), hence, selective attention was engaged as in a typical continuous performance attention task. In the second block, green rocket targets were frequent (67% of trials), hence, participants developed a prepotent impulse to respond. As individuals must intermittently suppress a motor response to sparse non-targets (33% of trials), this block provided a metric of response inhibition (Aron, 2007; Aron and Poldrack, 2005; Chambers et al., 2009).

##### 2. Working Memory

Participants accessed a game named *Lost Star* that is based on the standard visuo-spatial Sternberg task (Sternberg, 1966). Participants were presented a set of test objects (stars); they were instructed to maintain the visuo-spatial locations of the test objects in working memory for a 3 sec delay period, and then responded whether a probe object (star) was or was not located in the same place as one of objects in the original test set. We implemented this task at the threshold perceptual span for each individual, i.e. the number of star stimuli that the individual could correctly encode without any working memory delay. For this, a brief perceptual thresholding period preceded the main working memory task, allowing for equivalent perceptual load to be investigated across participants (Lavie et al., 2004). During thresholding, the set size of the test stars was progressively increased from 1-8 stars based on accurate performance; 4 trials were presented at each set size and 100% performance accuracy led to an increment in set size; <100% performance led to one 4-trial repeat of the same set size and any further inaccurate performance aborted the thresholding phase. The final set size at which 100% accuracy was obtained was designated as the individual’s perceptual threshold.

Post-thresholding, the working memory task consisted of 48 trials presented over 2 blocks (Lenartowicz et al. 2014). Each trial initiated with a central fixation ‘+’ for 500 msec followed by a 1 sec presentation of the test set of star objects located at various positions on the screen, then a 3 sec working memory delay period, followed by a single probe star object for 1 sec, and finally a response time window of up to 1 sec in which participants made a yes/no choice whether the probe star had a matching location to the previously presented test set. A happy/sad face emoticon was used to provide accuracy feedback for 200 msec followed by a 500 msec ITI. Summary accuracy was also shown between blocks. The total task duration was 6 min.

##### 3. Interference Processing

Participants accessed a game named *Middle Fish*, an adaptation of the Flanker task (Eriksen and Eriksen, 1974), which has been extensively used to study interfering distractor processing (Lavie, Hirst and Fockert, 2004; Shipstead, Harrison and Engle, 2012). Participants were instructed to respond to the direction of a centrally located target (middle fish) while ignoring all flanking distractor fish. On congruent trials the flanker fish faced the same direction as the central fish, while on incongruent trials they faced the opposite direction. A brief practice of 4-trials preceded the main task of 96 trials presented over two blocks for a total task time of 8 min. 50% of trials had congruent distractors and 50% were incongruent. To retain attention, the array of fish was randomly presented in the upper or lower visual field on equivalent number of trials. On each trial, a central fixation ‘+’ appeared for 500 msec followed by a 100 msec stimulus array of fish and up to a 1 sec response window in which participants responded left/right as per the direction of the middle fish. Subsequently a happy/sad face emoticon was presented for 200 msec for accuracy feedback followed by a 500 msec ITI. Summary accuracy was shown between blocks and the total task duration was 8 min.

##### 4. Emotional Interference Processing

We embedded this task in *BrainE* given ample evidence that emotions impact cognitive control processes (Gray, 2004; Pessoa, 2009; Inzlicht, Bartholow and Hirsh, 2015). Participants accessed a game named *Face Off*, adapted from prior studies of attention bias in emotional contexts (López-MartÍn *et al*., 2013, 2015; Thai, Taber-Thomas and Pérez-Edgar, 2016). We used a standardized set of culturally diverse faces from the Nim-Stim database for this assessment (Tottenham *et al*., 2009). We used an equivalent number of males and female faces, each face with four sets of emotions, either neutral, happy, sad or angry, presented on equivalent number of trials. An arrow was superimposed on the face on each trial, occurring either in the upper or lower central visual field on equal number of trials, and participants responded to the direction of the arrow (left/right). Participants completed 144 trials presented over three equipartitioned blocks with shuffled, but equivalent number of emotion trials in each block; a practice set of 4-trials preceded the main task. Each trial initiated with a central fixation ‘+’ for 500 msec followed by a face stimulus with a superimposed arrow of 300 msec duration. As in other tasks, participants responded within an ensuing 1 sec response window, followed by a happy/sad emoticon feedback for accuracy (200 msec) and a 500 msec ITI. Summary block accuracy feedback was provided, and the total task duration was 10 min.

#### Electroencephalography (EEG)

EEG data was collected simultaneous to all cognitive tasks using a 24-channel SMARTING device with a semi-dry and wireless electrode layout (Next EEG — new human interface, MBT). Data were acquired at 500 Hz sampling frequency at 24-bit resolution. Cognitive event markers were integrated using LSL and data files were stored in xdf format.

#### Repetitive Transcranial Magnetic Stimulation (rTMS)

In the second study, we used the FDA-approved Magventure stimulator (MagPro R30) for rTMS delivery. Each participant made two visits for this study, separated by a one-week interval, and each visit lasted up to 2 hours. Participants were provided either the continuous theta burst stimulation (cTBS) or intermittent TBS (iTBS) TMS protocol at each visit. Participants were blinded to the stimulation type, and stimulation order in week 1 or 2 was counterbalanced across subjects. The research staff who performed stimulation were blind to the effects of the cTBS or iTBS protocol, and the data analytics lead and study principal investigator were blind to the identity of the protocol i.e. all data were analyzed with cTBS blinded as stim A and iTBS as stim B. TBS stimulation was delivered to the midline at FCz target location, consistent with the pre-supplementary motor area site for rTMS in superior frontal cortex, which was active in most of our cognitive tasks (Verbruggen *et al*., 2010). A train of 3 pulses, spaced 20 msec apart (50 Hz stimulation), followed by an inter-train interval of at least 200 msec (5 Hz) was applied either continuously (cTBS), or intermittently (iTBS) with a jitter between trains as has been tested in prior research (Rossi, Hallett, Rossini, Pascual-Leone, *et al*., 2009; Oberman *et al*., 2011). In cTBS, bursts of 3 pulses at 50 Hz were applied at a frequency of 5 Hz for 20 sec, total 100 bursts. In iTBS, ten 2 sec periods (10 bursts) of TBS were applied at a rate of 0.1 Hz for a total 100 bursts. Stimulation amplitude was set at 80% of motor threshold individually determined in each participant.

At each rTMS study visit, participants first performed *BrainE* assessments (pre-stim), then immediately received either cTBS or iTBS TMS stimulation, then performed *BrainE* again (post-stim). This within subject test-retest method allowed us to test for reliability of *BrainE* assessment data, comparing pre-stim week 1 versus pre-stim week 2 results. Additionally, we investigated the sensitivity of BrainE assessments to measure brain plasticity in pre-stim versus post-stim comparisons, as a function of different cognitive operations and rTMS protocols. Figure 1F shows the rTMS study design.

### Data acquisition

**Participants**.102 adult human subjects (mean age 24.8 ± 6.7 years, range 18-50 years, 57 females) participated in the *BrainE* neuro-cognitive assessment study. Participants were recruited using IRB-approved on-campus flyers at UC San Diego as well as via the online recruitment forum, ResearchMatch.org, which hosts a registry of research volunteer participants; the ad on the Research Match registry was customized for participants in the general San Diego area (within 50 miles of our research location). Overall, ∼50% of participants were university affiliates (lab members and students), while the rest were from the general population (i.e., Research Match registry). All participants provided written informed consent for the study protocol (#180140) approved by the University of California San Diego institutional review board (UCSD IRB). Participant selection criteria included healthy adult status, i.e. without any current diagnosis for a neuropsychiatric disorder and/or current/recent history of psychotropic medications and/or hospitalization within the past 8 weeks. Five participants were excluded from the study as they had a current diagnosis for a psychiatric disorder and/or current/recent history of psychotropic medications. All participants reported normal/corrected-to-normal vision and hearing and no participant reported color blindness, majority of participants (95 of 102) were right handed. All participants had at least a high-school education (16 years). Unfortunately, we did not collect information on highest qualification.

For the two-visit TMS study, we enrolled 24 human subjects (mean age 24.3 ± 7.4 years, 17 females). 13 of these individuals had previously participated in the main *BrainE* assessment above, with a minimum one-month gap between participation in the two studies. Participants provided written informed consent for the TMS study protocol (#190059) approved by the UCSD IRB. The TMS study was pre-registered on Clinicaltrials.gov (NCT03946059). Participants were screened for this study prior to enrollment. Any individuals with a history of seizure disorder; vascular, traumatic, tumoral, infectious or metabolic lesion of the brain; administration of drugs that lower the seizure threshold; implanted or non-removable metallic objects above the neck; implanted devices with electrical circuits (pace-makers, cochlear implants) were excluded from enrollment. In addition, subjects were excluded if they had chronic sleep deprivation or confirmed heavy alcohol use (defined as greater than 5 episodes of binge drinking in the past month with >5 alcohol drink-equivalents per sitting for men (or >4 drink-equivalents per sitting for women). Subjects were also excluded if they reported the use of stimulant drugs in the past month (cocaine, methamphetamines), or if they were pregnant, or had any history of severe cardiovascular disease (i.e. history of transient ischemic attack, heart attack or stroke).

### Behavioral and Neural Processing Methods

#### Behavioral analyses

Behavioral data for all cognitive tasks were analyzed for signal detection sensitivity, d’, computed as z(Hits)-z(False Alarms) (Heeger and Landy, 2009). Task speeds were calculated as log(1/RT), where RT is response time in milliseconds. Task efficiency was calculated as a product of d’ and speed (Barlow *et al*., 1980; Vandierendonck, 2017). d’, speed, and efficiency metrics were checked for normal distributions prior to statistical analyses.

#### Neural Analyses

We applied a uniform processing pipeline to all EEG data acquired simultaneous to the cognitive tasks. This included: 1) data pre-processing, 2) computing event related spectral perturbations (ERSP) for all channels, and 3) cortical source localization of the EEG data filtered within relevant theta, alpha and beta frequency bands. 1)Data preprocessing was conducted using the EEGLAB toolbox in MATLAB (Delorme and Makeig, 2004). EEG data was resampled at 250 Hz, and filtered in the 1-45 Hz range to exclude ultraslow DC drifts at <1Hz and high-frequency noise produced by muscle movements and external electrical sources at >45Hz. We performed 827-point bandpass, zero phase, filtering with transition band width 4.063Hz, and passband edges of [1 45] Hz for cleaning the epoched data of time length [-1.5 1.5] secs; [3 7] Hz for theta specific filtering, [8 12] Hz for alpha specific and [13 30] Hz for beta specific data analysis. EEG data were average referenced and epoched to relevant stimuli in each task, as informed by the LSL time-stamps. While 24 channels is not a dense set, they are far enough from each other that no common neural signature is removed, but only common in-phase noise present in all channels is canceled during average referencing (Nunez, 2010). Any task data with missing LSL markers (1.4% of all data) had to be excluded from neural analyses. Any missing channel data (channel F8 in 2 participants) was spherically interpolated to nearest neighbors. Epoched data were cleaned using the autorej function in EEGLAB to remove noisy trials (>5sd outliers rejected over max 8 iterations; 6.6± 3.4% of trials rejected per participant). EEG data were further cleaned by excluding signals estimated to be originating from non-brain sources, such as electrooculographic, electromyographic or unknown sources, using the Sparse Bayesian learning (SBL) algorithm (Ojeda et al., 2018, 2019, https://github.com/aojeda/PEB) explained below.

2) For ERSP calculations, we performed time-frequency decomposition of the epoched data using the continuous wavelet transform (cwt) function in MATLAB’s signal processing toolbox. Baseline time-frequency (TF) data in the −750 msec to −550 msec time window prior to stimulus presentation were subtracted from the epoched trials (at each frequency) to observe the event-related synchronization (ERS) and event-related desynchronization (ERD) modulations (Pfurtscheller, 1999).

3) Cortical source localization was performed to map the underlying neural source activations for the ERSPs using the block-Sparse Bayesian learning (SBL) algorithm (Ojeda, Kreutz-Delgado and Mullen, 2018; Ojeda *et al*., 2019) implemented in a recursive fashion. This is a two-step algorithm in which the first-step is equivalent to low-resolution electromagnetic tomography (LORETA, (Pascual-Marqui, Michel and Lehmann, 1994). LORETA estimates sources subject to smoothness constraints, i.e. nearby sources tend to be co-activated, which may produce source estimates with a high number of false positives that are not biologically plausible. To guard against this, SBL applies sparsity constraints in the second step wherein blocks of irrelevant sources are pruned. Source space activity signals were estimated and then their root mean squares were partitioned into 1) regions of interest (ROIs) based on the standard 68 brain region Desikan-Killiany atlas (Desikan et al. 2006; **Supplementary Figure 1**) using the Colin-27 head model (Holmes *et al*., 1998) and 2) artifact sources contributing to EEG noise from non-brain sources such as electrooculographic, electromyographic or unknown sources; activations from non-brain sources were removed to clean the EEG data. The SBL GUI accessible through EEGLAB provides access to an EEG artifact dictionary; this dictionary is composed of artifact scalp projections and was generated based on 6774 ICs available from running Infomax ICA on two independent open-access studies (http://bnci-horizon-2020.eu/database/data-sets, study id: 005-2015 and 013-2015). The k-means method is used to cluster the IC scalp projections into Brain, EOG, EMG, and Unknown components. We checked visually that EOG and EMG components had the expected temporal and spectral signatures according to the literature (Jung et al., 2000). The SBL algorithm returns cleaned channel space EEG signals in addition to the derived cortical source signals as outputs. In this study, we first applied SBL to the epoched channel EEG signals; activations from artifact sources contributing to EEG noise, i.e., from non-brain sources such as electrooculographic, electromyographic or unknown sources, were removed to clean the EEG data (Ojeda *et al*., 2019). Cleaned subject-wise trial-averaged channel EEG data were then specifically filtered in theta (3-7 Hz), alpha (8-12 Hz), and beta (13-30 Hz) bands and separately source localized in each of the three frequency bands and in each task to estimate their cortical ROI source signals. The source signal envelopes were computed in MatLab (envelop function) by a spline interpolation over the local maxima separated by at least one time sample; we used this spectral amplitude signal for all neural analyses presented here. We focused on post-stimulus encoding in the 100-300 msec range for theta and alpha bands, and 400-600 msec spectral amplitude range for the beta band signals, respectively. These epoch windows were chosen based on the peak global activity of the task-averaged signals in the respective frequency bands. We used these time windows to compute common-task-average neural signals and also distinct-task based neural activations across subjects.

### Statistical Analyses

Behavioral data were compared across tasks using repeated measures analyses of variance (rm-ANOVA) with a within-subject factor of task-type; the Tukey-Kramer method was used for post-hoc testing.

Channel-wise theta, alpha, beta ERS and ERD modulations on each task, and within the common-task-average were analyzed for significance relative to baseline using t-tests (p≤0.05), followed by false discovery rate (fdr) corrections applied across the three dimensions of time, frequency, and channels (Genovese, Lazar and Nichols, 2002).

Significant source activations underlying the theta, alpha, beta ERS and ERD modulations were computed using t-tests with Bonferroni family wise error rate (fwer) correction applied for multiple comparisons in the 68 ROI source dimension, 5 tasks and 3 frequency bands (p≤0.00005). For the global cognitive task-averaged activity averaged across 5 tasks, the modulations were computed using t-tests fwer correction applied for multiple comparisons in the 68 ROI source dimension and 3 frequency bands (p≤0. 00024). Rm-ANOVA tests were conducted to investigate differences in frequency band x task type cortical activations, and the Tukey-Kramer method was used for post-hoc tests.

For the TMS study, we first calculated the Cronbach’s alpha internal consistency measure (MatLab Intraclass Correlation Coefficient, ICC, type ‘C-k’ function) for the week 1 vs week 2 pre-data to assess reliability of the cognitive performance metrics as well as neural signals at each cortical source region. Additionally, we conducted rm-ANOVA tests with within-subjects factors of stimulation type (cTBS vs iTBS) and assessment time (pre-vs post-); results were corrected for multiple comparisons across 5 cognitive tasks and 3 frequency bands at p≤0.003 significance threshold, the significant ROIs were further corrected for multiple comparisons using fdr; the Tukey-Kramer method was used for post-hoc tests. Estimates of effect size were calculated as standardized mean difference/Cohen’s d (Cohen *et al*., 1988) with the Hedges and Olkin small sample bias correction applied (Hedges and Olkin, 1985).

Finally, we investigated the relationship between the cognitive and neural activations versus subjective mental health symptom severity for anxiety, depression, inattention and hyperactivity self-reports using Spearman correlations (thresholded at p≤0.05). For neural data, we used the significant global cognitive task-average activity for correlations. For the four symptom data that were highly correlated, we conducted a principal component analysis (PCA) and used the top PC that explained majority of the symptom score variance across subjects, and further corrected for multiple comparisons across ROIs and frequency bands using fdr. We confirmed Spearman correlations were appropriate for correlations based on the Anderson-Darling test for normality (Spearman, 1904; Anderson *et al*., 1952) and confidence intervals were calculated using 10,000-iteration percentile bootstrap method (Efron, 1982).

## Results

### Behavioral performance

Signal detection sensitivity d’, response times (msec), speed and efficiency for all tasks are shown in Table 1. Repeated measures ANOVAs were conducted on each behavioral variable with five task types as within-subjects factor. For d’, we found a significant effect of task (F_4,384_ = 218.22, p<0.0001). Post-hoc tests revealed significant interactions between every task type pair (p<0.05). For speed and efficiency, we again found a significant effect of task (speed: F_4,384_ = 559.29, p<0.0001; efficiency: F_4,384_ = 715.13, p<0.0001) and the post-hoc tests for each of them showed significant interaction between each task type pair (p<0.001) except for that between interference processing and emotion interference processing tasks for speed.

**Table 1.** Behavioral performance across tasks for all participants (n=97), as mean ± standard error of mean (sem). Response times that did not have a normal distribution, are reported as median ± median absolute deviation (mad).

| Cognitive Task | *d'*<br>mean ± std | Response time<br>median ± mad sec | Speed<br>mean ± std | Efficiency<br>mean ± std |
| --- | --- | --- | --- | --- |
| Selective attention | 4.47 ± 0.32 | 0.44 ± 0.03 | 0.36 ± 0.05 | 0.34 ± 0.06 |
| Response inhibition | 4.28 ± 0.46 | 0.40 ± 0.04 | 0.40 ± 0.06 | 0.36 ± 0.07 |
| Working memory | 2.06 ± 0.92 | 0.88 ± 0.14 | 0.04 ± 0.10 | 0.02 ± 0.05 |
| Interference processing | 3.63 ± 0.83 | 0.48 ± 0.03 | 0.31 ± 0.05 | 0.24 ± 0.06 |
| Emotion interference processing | 3.38 ± 0.65 | 0.48 ± 0.03 | 0.31 ± 0.06 | 0.22 ± 0.05 |

### Neural activations at EEG channels

Results of the time-frequency decompositions of the stimulus-evoked neural activity are shown at exemplar electrodes, FCz and POz, for all five tasks and for the global cognitive average across tasks (**Figure 2**). ERS/ERD modulations in the data were fdr-corrected across time, frequency and channel dimensions across subjects. Most tasks had significant and equivalent ERS and ERD signatures at the channel level, with ERS predominant in the theta/alpha frequencies and ERD predominant in the beta frequency range. We also show topographic maps in each task (Figure 2) for the stimulus-evoked peak activity windows and for frequency averaged theta, alpha, beta, during the 100-300 msec time range for theta and alpha, and 400-600 msec range for beta.

**Figure 2.**
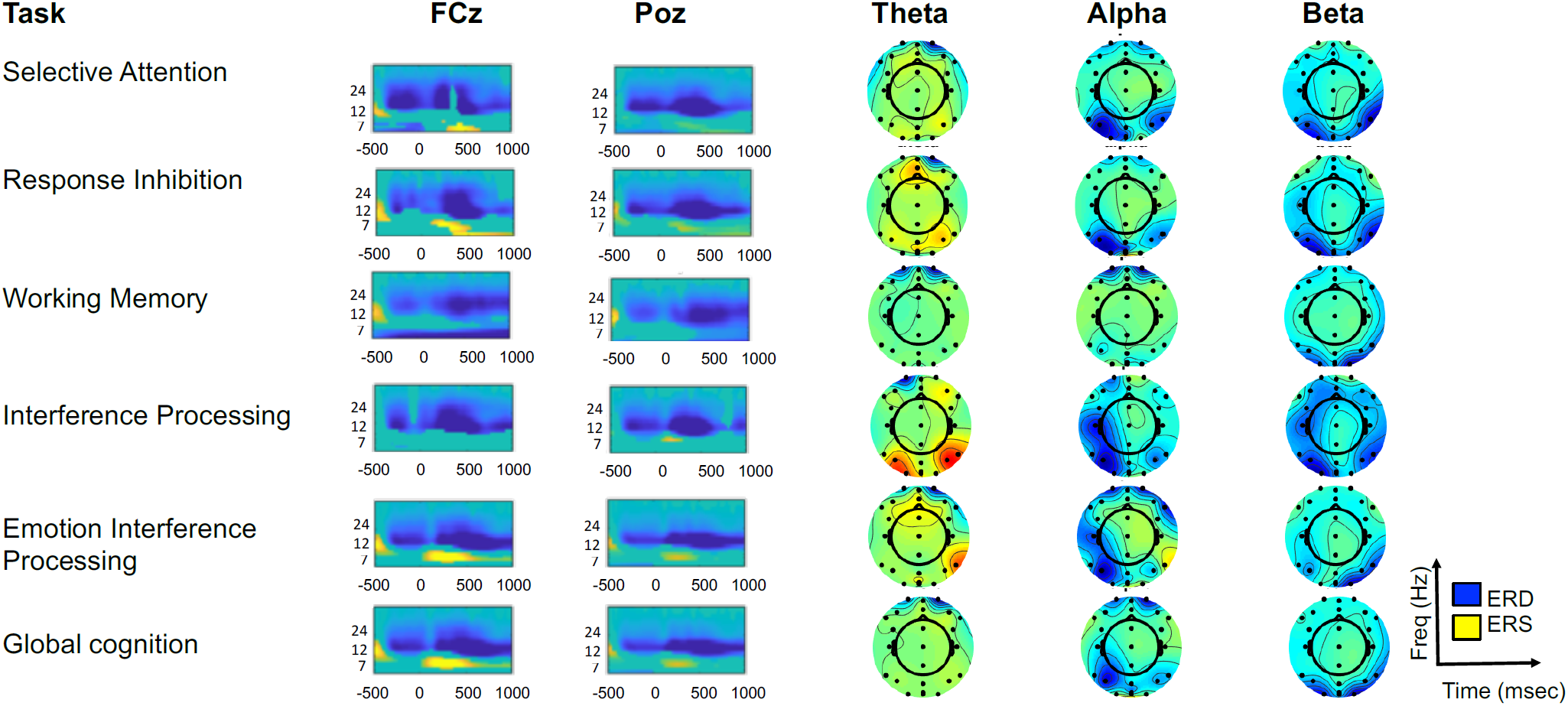
Significant event related synchronization (ERS: yellow) and desynchronization (ERD: navy blue), fdr-corrected, time-locked to the stimulus (−0.5 to +1 sec) across all tasks and averaged across tasks (global cognition) at exemplar electrodes FCz and POz. ERS was observed at theta/alpha frequencies while ERD was predominant in the beta frequency range. Topographic maps for the stimulus-evoked peak activity windows for the frequency-averaged theta, alpha and beta band signals are shown at right, for the peak time windows, of 100-300 msecs for theta and alpha, and 400-600 msecs for beta.

### Neural activations at cortical sources

Significant cortical source-localized neural activity in the theta, alpha and beta bands for the stimulus encoding period, for each cognitive task and for the global task-average are shown in **Figure 3**; both p<0.05 uncorrected and p≤0.00005 fwer-corrected maps for individual tasks, p≤0.00024 fwer-corrected global cognition maps are shown. Consistent with the channel maps, theta and alpha frequencies predominantly showed ERS at bilateral cortical sites, while significant ERD was observed for beta frequencies prominently in medial frontal, parietal, posterior cingulate cortex and left sensorimotor cortex.

**Figure 3.**
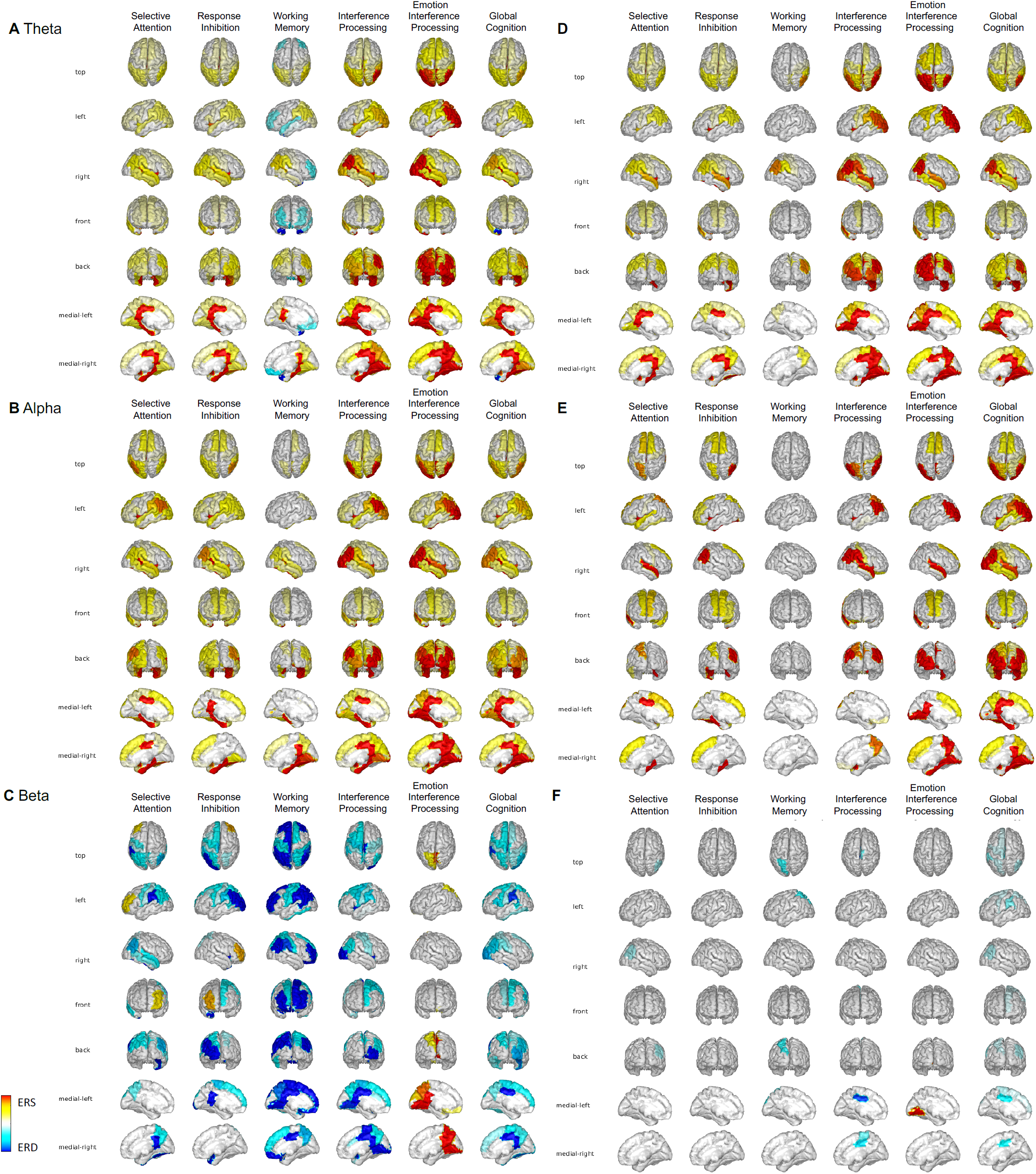
Significant theta, alpha and beta band ERS and ERD signatures during stimulus encoding relative to baseline for the five cognitive tasks and for the global cognitive task-average. Left column maps are at uncorrected p<0.05 threshold and right column maps are fwer corrected.

We conducted repeated measured ANOVAs on the source activations at the 68 ROIs with the three frequency bands and five task types as within-model factors, to investigate whether theta, alpha and beta band modulation patterns in cortical space were significantly different from each other across tasks. These analyses showed a main effect of frequency band (F_2,134_=13.65, p<0.0001) and task (F_4,268_ = 12.79, p<0.0001) and a significant frequency band x task interaction (F_8,536_=10.40, p<0.0001). Post-hoc tests revealed significant differences in the three frequency band cortical activations in each task (p<0.01) except in the working memory task where there were no significant differences in theta/alpha/beta specific cortical maps.

Overall, the theta and alpha band cortical activations were significantly positively correlated, while these lower frequency activations were negatively correlated with the beta band activations (Pearson correlations on the global cognitive task-average data (p<0.01); **Supplementary Figure 2**). We also confirmed that the cortical activations maps were near equivalent if they were computed over all task trials vs. just correct trials (93.50±3.45% correct trials averaged across tasks). These all vs. correct trial maps were strongly positively correlated in each frequency band (Spearman correlations on the global cognitive task-average maps across 68 ROIs, r(67)>0.99, p<0.001 for theta and alpha bands and r(67)=0.78, p<0.001 for beta band).

### Common and distinct neural activations across cognitive tasks

We computed logical maps representing whether each cortical source was significantly active in one or more tasks regardless of the neural activity magnitude (binarized at p≤0.00005 fwer threshold, Figure 4A); if any cortical source ROI was uniquely active in a single cognitive task, we further identified that distinct task (Figure 4B). These maps showed neural activations in brain-wide ROIs in the theta/alpha bands in the majority of tasks, with greater cortical overlap in the theta than alpha band (up to 3 tasks). In the beta band there was no overlapping task activity.

**Figure 4.**
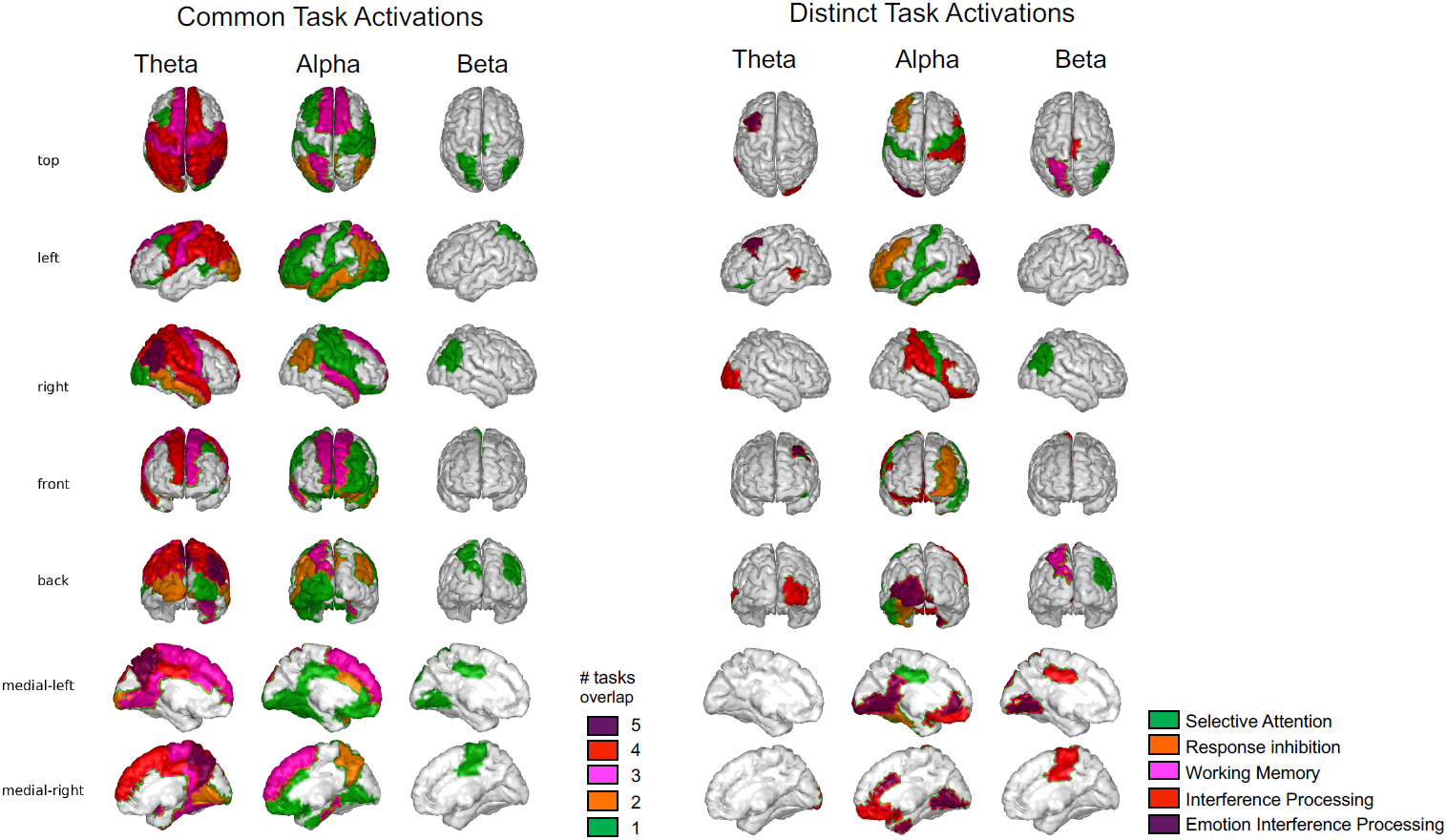
Common and distinct neural activations across tasks. (A) Common activation brain maps are logical maps showing cortical sources that are active during stimulus encoding in one or more cognitive tasks. (B) Distinct activation brain maps are logical maps showing cortical sources that are active during stimulus encoding only in one particular cognitive task. Logical maps are based on significantly active ROIs within task at p≤0.00005 fwer threshold.

Distinct task activation maps revealed that only the selective attention task significantly activated left inferior frontal cortex, bilateral sensory-motor cortices, left superior temporal cortices in the alpha band. The response inhibition task selectively activated the left caudal middle frontal area, commonly referred to as dorsolateral prefrontal cortex (DLPFC), in the alpha band. Activity related to stimulus-encoding on the working memory task was observed in left superior parietal cortex in the beta band. The Flanker interference task selectively activated right inferior frontal cortex, bilateral orbitofrontal, right sensory and supramarginal cortices in the alpha band, bilateral posterior cingulate in the beta band. Finally, the emotion interference task selectively activated left caudal middle frontal area/dlPFC in the theta band as well as left orbitofrontal, right anterior cingulate, left posterior/isthmus-cingulate cortex and bilateral fusiform regions in the alpha band, and left fusiform activation in the beta band, potentially specific to face stimuli in this task.

### TMS driven cognitive neuroplasticity

In this second study, participants made two visits completing the five *BrainE* cognitive tasks twice at each visit, pre- and post-rTMS application. Either the cTBS or iTBS protocol for rTMS was applied at each visit counterbalanced across subjects (see Methods, Figure 1F). We calculated Cronbach’s alpha as a summary reliability measure for the pre-stim visit 1 vs. visit 2 cognitive and neural data. For cognitive performance across the 25 healthy subjects, reliability was high (task-averaged Cronbach’s alpha for d’: 0.83, speed: 0.80, efficiency: 0.80, p<0.0001). For neural activity averaged across all five tasks and concatenated across all three frequency bands and summarized across cortical source sites, reliability was high (Cronbach’s alpha = 0.77, p<0.0001). When neural data were further analyzed separately for reliability in the three frequency bands, Cronbach’s alpha values were more variable (for theta: 0.55, p<0.05; alpha: 0.73, p=0.001; beta: 0.44, p=0.08), though paired t-tests confirmed that visit 1 vs. 2 pre-stim neural data were not significantly different in any frequency band (all p>0.05). Finally, we also calculated neural reliability concatenated across all three frequency bands within each cortical source region, showing moderate to high test-retest reliability across different cortical ROIs (Figure 5A).

**Figure 5.**
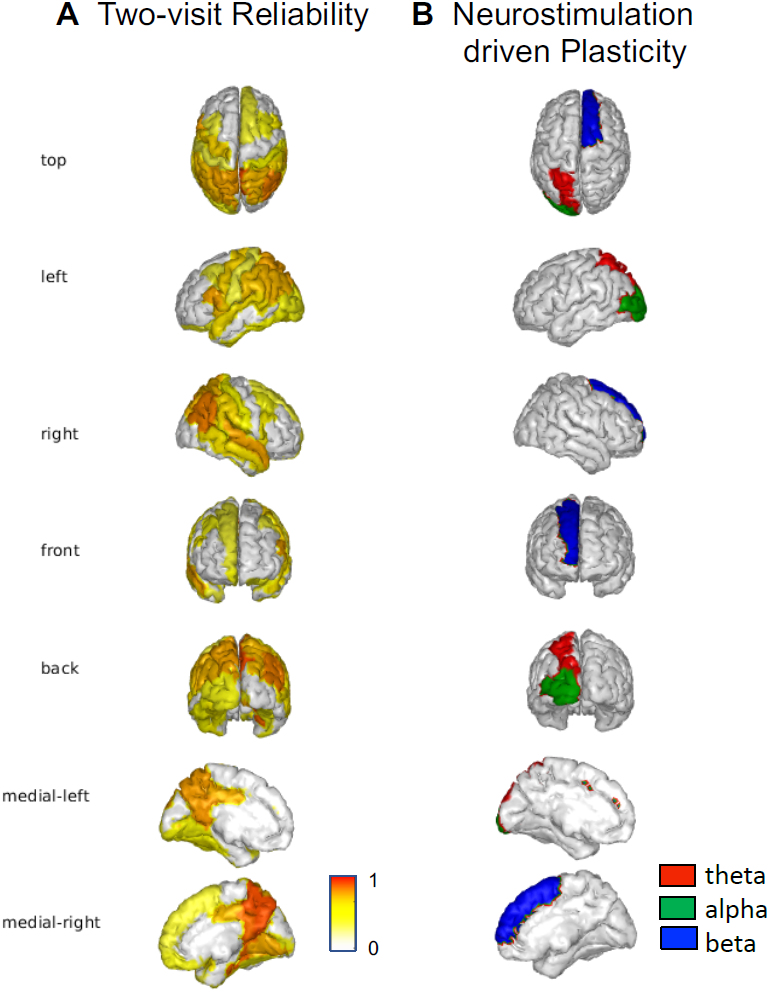
TMS study results. (A) Neural data acquired at pre-TMS at visit 1 and 2 showed moderate to high reliability measured across all three frequency bands using Cronbach’s alpha calculated for each cortical source region [min: 0.5 max: 0.9, thresholded at p<0.05]. (B) Significant stimulation type (iTBS vs. cTBS) by time (pre-vs. post-stim) neural interactions in rm-ANOVAs emerged only for the emotion interference processing task, shown in red for theta band, green for alpha band and blue for beta band activations (post-pre iTBS>cTBS, p≤0.003 fwer correction applied for 5 tasks x 3 frequency comparisons and fdr-corrected for brain-wide ROI multiple comparisons).

We performed a 3-factor repeated measures ANOVA for each behavioral measure (d’, speed, efficiency) with task type, assessment time (pre-stim, post-stim) and stimulation type (cTBS, iTBS) as within-subject factors. No analyses showed main effects or interactions for stimulation type, thus single session rTMS stimulation did not affect cognitive behaviors. A significant main effect of task type was found for each behavioral measure (d’: F_4,92_=188.47.65, p<0.0001; speed: F_4,92_=159.17, p<0.0001; efficiency: F_4,92_=342.34 p<0.0001), and similarly assessment time also showed a significant main effect for each measure (d’: F_1,23_=6.53, p=0.02; speed: F_1,23_=81.27, p<0.0001; efficiency: F_1,23_=38.42, p<0.0001); a significant task x assessment time interaction only emerged for the speed measure (F_4,92_=9.86, p<0.0001) that showed significantly greater speed at post vs. pre for all tasks (p<0.01). Post-hoc pre/post speed comparisons (Tukey-Kramer test) showed a larger post vs. pre change for working memory (Δspeed, 0.04±0.006, p<0.0001) followed by that for sustained attention (Δspeed, 0.02±0.004, p=0.0009) and emotion interference processing (Δspeed, 0.02±0.003, p<0.0001) and then for flanker interference (Δspeed, 0.01±0.003, p=0.008) and response inhibition (Δspeed, 0.01±0.004, p=0.003), but with no differential effect by stimulation type.

In rm-ANOVAs conducted on the neural data, we explicitly focused on significant stimulation type x assessment time interactions to understand differential neuroplasticity outcomes of cTBS vs iTBS. Results were thresholded at p≤0.003 fwer for 5 tasks x 3 frequency band comparisons and fdr-corrected for multiple comparisons across all ROIs (Figure 5B). These interactions exclusively showed significance for the emotional interference task in the left superior parietal brain region with a large effect size in the theta band (Cohen’s d, iTBS>cTBS, 1.32, 95% CI [0.7 1.94], p=0.0011), in left lateral occipital area with a medium effect size in the alpha band (Cohen’s d, iTBS>cTBS, 0.65, 95% CI [0.07 1.23], p=0.0006), and in right superior frontal/rostral anterior cingulate cortex with a large effect size in the beta band (Cohen’s d, iTBS>cTBS, 1.09, 95% CI [0.49 1.70], p=0.0029).

### Cognitive neural correlates of subjective mental health

All participants provided self-reports on standard scales of anxiety, depression, inattention and hyperactivity. These four symptoms had high inter-correlation coefficients (mean±sem, r= 0.57±0.03, p<0.0001) in our participant sample. Hence, we conducted a PCA of the symptoms and extracted the top mental health PC that explained 69.72% variance in the symptom data; other PC components were not considered as they each explained less than a quarter of the total variance. Spearman correlations of the cognitive metrics (d’/speed/efficiency) with the mental health PC did not show any significant correlations (all p>0.05).

For mental health correlations with neural data, we focused on the significant global task-averaged evoked activity (Figure 3, rightmost column) and found several symptom correlates, specifically in the theta and alpha bands (**Table 2** and **Figure 6**, p<0.05 corrected for multiple comparisons across ROIs and frequency bands using fdr, associated scatter plots shown in **Supplementary Figure 3**). In all cases, more severe symptoms across our healthy participant sample were associated with significantly reduced ERS activity. Theta/alpha symptom correlates were widespread and included distinct cognitive control regions of the fronto-parietal network including the left DLPFC (Caudal middle frontal L in Table 2), temporal regions, and visual areas such as the precuneus showed negative correlations with the symptom PC.

**Table 2.**
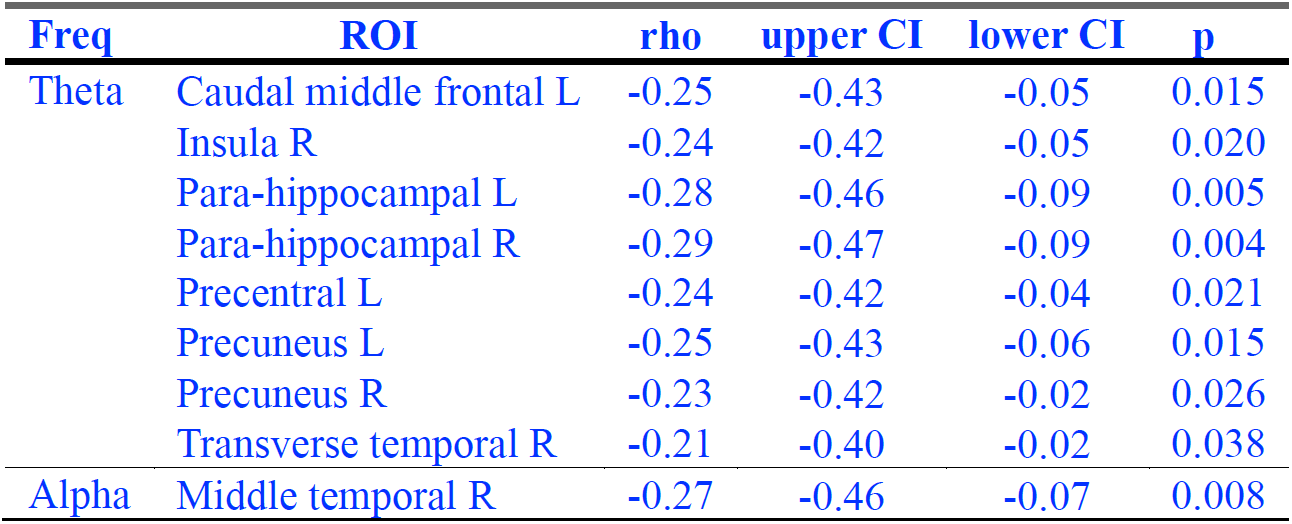
Global task-averaged neural correlates of subjective mental health symptoms. Significant correlates were observed in theta and alpha frequency bands (Spearman correlations, p < 0.05 fdr-corrected for multiple comparison across ROIs and frequency bands). Correlation coefficients, upper and lower 95% confidence intervals (CI) and p-values are shown

**Figure 6.**
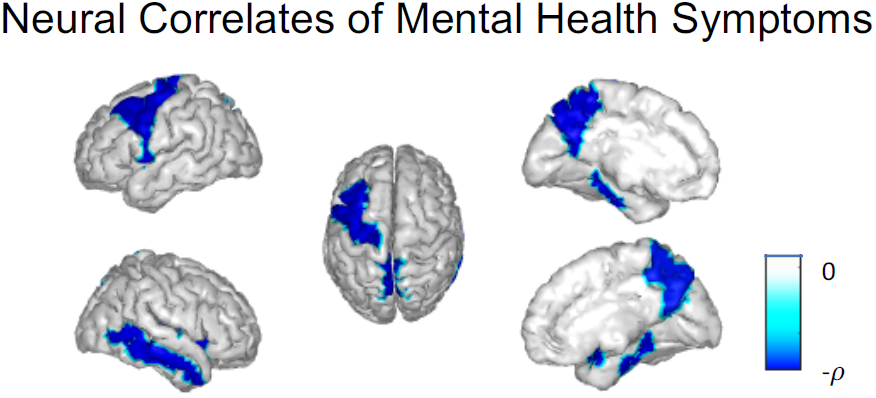
Global task-average neural correlates of subjective mental health. We found significant neural correlations with the top principal mental health symptom component that explained 69.72% variance across all symptoms. All correlations were negative and predominantly in the theta band (Table 2).

Correlations used the Spearman method (p<0.05, rho (*p*) values were fdr-corrected for multiple comparisons across ROIs and frequency bands).

## Discussion

In this study, we developed a scalable and accessible, mobile EEG based platform to assess neuro-cognitive processing. We refer to this platform as *BrainE* (short for Brain Engagement) and demonstrate how it can be used to inform cognitive and neural processes, specifically stimulus encoding, in five cognitive task contexts - selective attention, response inhibition, interference processing, emotional interference processing, and working memory. We present cortical processes parsed in distinct theta, alpha and beta frequency bands, with robust ERS in theta and alpha and inversely correlated ERD in the beta band. We demonstrate that the five tasks elicit common stimulus encoding related neural processing as well as some distinct cortical activations. In a second experiment that used rTMS to engender neuroplasticity, we show that specific cognitive tasks exhibit differential neural outcomes to two different, continuous versus intermittent, theta burst stimulation protocols.

Notably, we conducted these experiments with mobile wireless EEG in a rapid test sequence of less than an hour per subject. The results processed in the cortical space demonstrate consistency with findings from neuroimaging studies using less scalable approaches such as fMRI or high density EEG. Electrophysiological studies in primates and humans have shown that theta and alpha band evoked responses are broadly distributed in the brain and reflect many top-down cognitive control operations including goal directed attention, memory encoding, and novelty detection among others, consistent with our results (Aftanas and Golocheikine, 2001; Makeig *et al*., 2002; Ekstrom *et al*., 2005; Christie and Tata, 2009; Mishra *et al*., 2012; Buzsáki and Moser, 2013). In beta band global cognitive activity, we observed left lateralized sensory-motor ERD, contralateral to the right-hand dominant responses made by our participants, which is consistent with motor performance representations in prior studies (Aron, 2007; Picazio *et al*., 2014; Zavala, Jang, Trotta, Codrin I. Lungu, *et al*., 2018; Khanna and Carmena, 2017). Amongst some interesting activations, the response inhibition task specifically showed alpha ERS in left caudal middle frontal cortex (DLPFC area) and the selective attention task showed significant ERS in left inferior frontal cortex, which aligns with previous findings (Aftanas and Golocheikine, 2001; Palva and Palva, 2007; Zavala, Jang, Trotta, Codrin I Lungu, *et al*., 2018; Beltrán *et al*., 2019; Chong, Williams, Cunnington, and Mattingley, 2008). In our implementation, these two tasks only differed in the presented frequency of target vs. non-target stimuli and both tasks required moment-to-moment flexible decision-making whether to respond or to inhibit response; many studies show that DLPFC is important for such flexible decision-making (Dosenbach *et al*., 2007; Menon and Uddin, 2010) with noted alpha band oscillatory effects found in this region (Sadaghiani *et al*., 2012, 2019). The Flanker interference processing task significantly activated right inferior frontal cortex as well as right sensory-motor areas in the alpha band, which is in line with studies of interference control and inhibitory processing (Brass *et al*., 2005; Tettamanti *et al*., 2008; Hampshire *et al*., 2010; Zanto *et al*., 2011; Mishra *et al*., 2014; Zavala, Jang, Trotta, Codrin I Lungu, *et al*., 2018; Beltrán *et al*., 2019). The emotional interference processing task particularly activated left caudal middle frontal area or DLPFC in the theta band aligned with other neuroimaging studies using emotion tasks (Siegle *et al*., 2007; Grimm *et al*., 2008; Avissar *et al*., 2017); posterior (isthmus-) cingulate cortex in the alpha band was also modulated in this task as observed by others (Waugh, Lemus and Gotlib, 2014; Okon-Singer *et al*., 2015; Song *et al*., 2017). Finally, during working memory encoding, we observed distinct activity in the beta band that localized to parietal cortex matching prior evidence, especially with respect to right hemispheric activations (Berryhill and Olson, 2008; Nee *et al*., 2013). These results provide confidence that a mobile EEG tool, which can be easily scaled to any lab/community setting with limited resources, can be used to generate neuro-cognitive results that replicate the literature.

In the rTMS study, we first demonstrated that the task-related cognitive performance and neural processing data were reliably replicable across two baseline sessions completed one-week apart. Specifically, we computed intraclass correlation coefficients between the two baseline sessions that showed moderate-to-high reliability, particularly in the visual, parietal and temporal regions relative to the frontal activations, consistent with findings in other studies (McEvoy, 2000; Gudmundsson *et al*., 2007). We compared cognitive and neural effects of continuous (cTBS) and intermittent (iTBS) theta burst stimulation protocols, as previous studies have suggested their contrasting effects—iTBS to facilitate while cTBS to inhibit cortical excitability (Thimm and Funke, 2015; Viejo-Sobera *et al*., 2017; Vékony *et al*., 2018). No such differential effects were found for the cognitive performance measures; both stimulation protocols speeded up information processing as evidenced in post-vs. pre-stimulation response time differences in several tasks, most prominently on the working memory task. This was an interesting finding given that our healthy participant sample was already performing at high accuracy on the cognitive tasks, and suggests that rTMS application generally enhanced alertness (Guse, Falkai and Wobrock, 2010; Mensen *et al*., 2014). Absence of a sham rTMS arm limits further interpretation. Notably, medium to large effect size differential neural outcomes were observed for iTBS versus cTBS, particularly in the emotion interference processing task in all theta/alpha/beta frequency bands. The majority of these effects showed greater positive neuroplasticity for iTBS versus cTBS (W, 2005; Hoy *et al*., 2016). Modulations were observed in occipito-parietal brain regions in theta/alpha and in cognitive control regions of superior frontal/rostral anterior cingulate cortex in the beta band. The specificity of these results to certain tasks, brain regions and neural rhythms shows that the *BrainE* platform has utility for assessing rTMS related neuro-cognitive plasticity in future studies. Interestingly, rTMS is an FDA-approved treatment for depression (Rossi, Hallett, Rossini and Pascual-Leone, 2009; George, Taylor and Short, 2013), a disorder with emotion dysregulation problems. That we find neural processes on an emotion interference processing task sensitive to rTMS protocols suggests that this task could serve as a promising assay for measuring neuro-cognitive outcomes in future rTMS studies of depression.

While we investigated neuro-cognitive outcomes in healthy subjects excluding data from those with a clinical diagnosis, the study participants reported varying degrees of severity of anxiety, depression, inattention and hyperactivity symptoms. Self-reports were highly correlated across the four symptom scales, hence, we extracted the top principal component of the mental health symptoms. We found widespread mental health correlations of global task-averaged neural activity in the theta band, and a few activations in the alpha band. All correlations were negative showing reduced theta/alpha activity with greater symptom severity. The DLPFC/caudal middle frontal region, insular cortex were prominent in these neuro-behavioral correlations, aligned with studies demonstrating dysfunction in the core cognitive control networks, the fronto-parietal network and the cingulo-opercular network, in mood disorders (McNaughton, 1997; Deckersbach, Dougherty and Rauch, 2006; Zhao *et al*., 2007; Canbeyli, 2010; Brzezicka, 2013; Etkin, Gyurak and O’Hara, 2013) and in ADHD (Hesslinger *et al*., 2002; Biederman *et al*., 2008; Bush, 2011). Finally, we also found negative symptom correlations in the memory-related middle temporal area, and orbital network including para-hippocampal regions (Haldane and Frangou, 2006; Price and Drevets, 2012).

Overall, our research shows that the *BrainE* platform can serve as a useful tool to map several dimensions of neuro-cognition in a rapid, scalable and cost-effective manner. We further demonstrate that the tool can be used to study neuroplasticity of targeted interventions. In this study, the emotion interference processing task was most sensitive to differential neurostimulation protocols. We also show meaningful correlates of mental health symptoms. In future, this research platform can serve to inform the Research Domain Criteria (RDoc) framework for investigating mechanisms of mental disorders (Insel *et al*., 2010), both in terms of understanding the neuro-cognitive correlates of mental disorders and to study specific circuit engagement in the context of targeted interventions that engage neuroplasticity. Notably, we quantify several analyses in cortical source space, thus, facilitating comparison with the EEG as well as fMRI literature. While this particular study was limited to a healthy adult cohort, we aim to integrate this platform in future neuro-cognitive studies in children and adolescents, aging adults, as well as individuals with clinical psychiatric diagnoses. Given the mobility of the *BrainE* platform, it is not limited to the research lab setting, and can be used to reach participants and acquire data in community settings such as schools and clinics, enabling greater diversity in research participation (Mishra, 2019). Finally, we have only scratched the surface of the rich neural dynamics that can be investigated in this dataset, limiting the neural analyses in this study to stimulus-evoked spectrotemporal activity modulations on core cognitive tasks; future studies may investigate aspects of functional connectivity as well as information processing in the context of task cues, and onset of responses and rewards on these tasks and newly added cognitive tasks. Fundamentally, the *BrainE* platform enables systematic cognitive neuroscience studies at scale across the mental health spectrum. In future, it may be used to find new biomarkers of brain-targeted interventions, and its ease of use may help to reduce the replicability crisis of small sample lab studies.

### CRediT author statement

Pragathi Priyadharsini Balasubramani: Conceptualization, Methodology, Formal Analysis and Investigation, Writing-Reviewing and Editing

Alejandro Ojeda: Methodology, Writing

Gillian Grennan, Vojislav Maric, Hortense Le, Fahad Alim, Mariam Zafar-Khan, Juan Diaz-Delgado, Sarita Silveira: Investigation and Data Curation

Dhakshin Ramanathan: Conceptualization, Methodology

Jyoti Mishra: Conceptualization, Methodology, Reviewing and Editing, Supervision

## Acknowledgements

This work was supported by University of California San Diego (UCSD) lab start-up funds (DR, JM), the Interdisciplinary Research Fellowship in NeuroAIDS (PB: R25MH081482), the Burroughs Wellcome Fund Career Award for Medical Scientists (DR) and the VA Medical Center Career Development Award (DR: 7IK2BX003308). We thank Alankar Misra for software development of the *BrainE* software and several UCSD undergraduate students who assisted with data collection. The *BrainE* software is copyrighted for commercial use (Regents of the University of California Copyright #SD2018-816) and free for research and educational purposes.

## Data Availability

The dataset in this study is available on the open-access repository link: 10.5281/zenodo.4088951

---

**Supplementary Figure 1.**
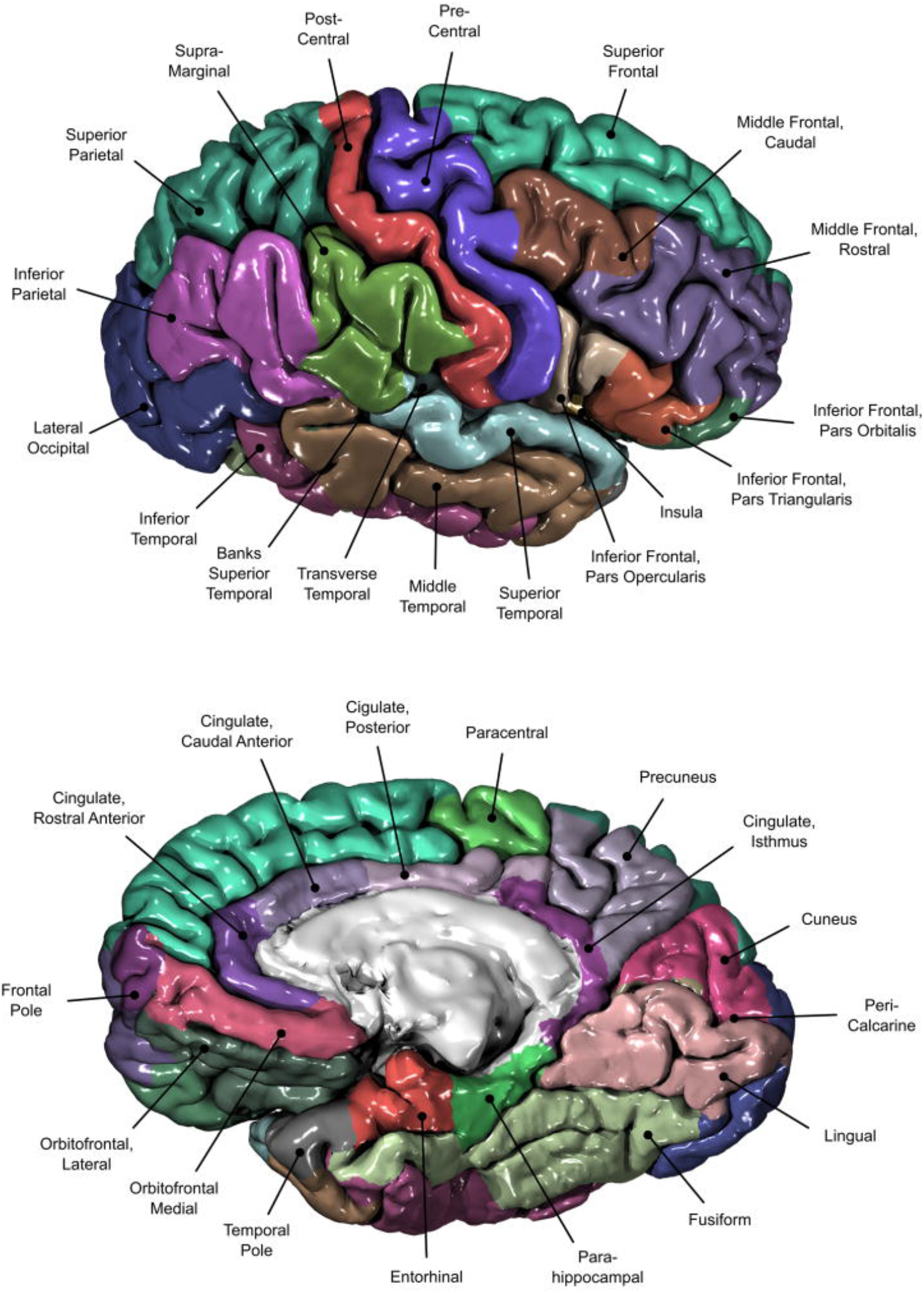
Cortical source regions as per the Desikan-Killiany atlas (Desikan *et al*., 2006).

**Supplementary Figure 2.**
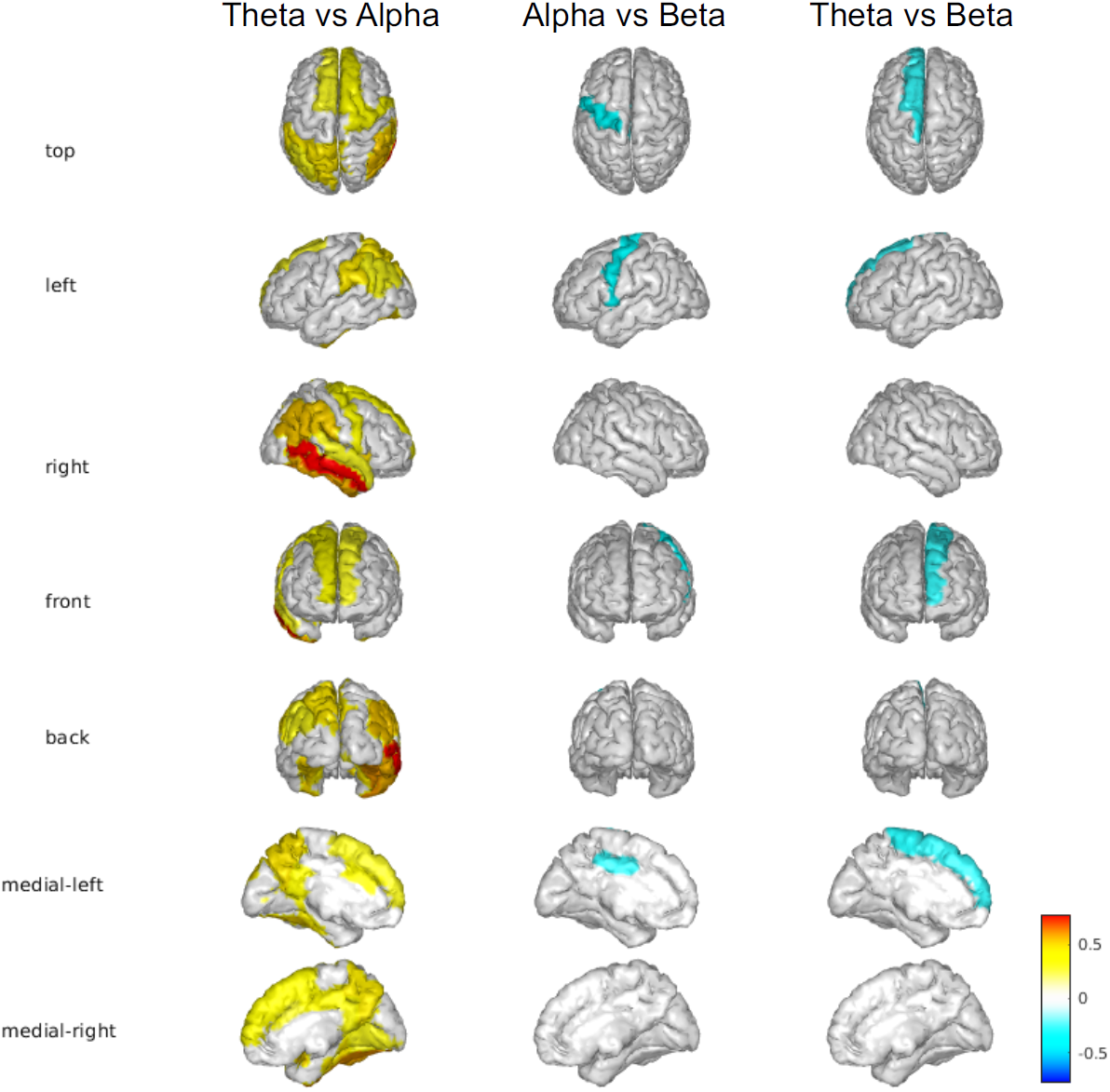
Significant correlations between the theta, alpha and beta band global cognitive task-average maps are shown (Pearson correlations, r(96) at p<0.01).

**Supplementary Figure 3.**
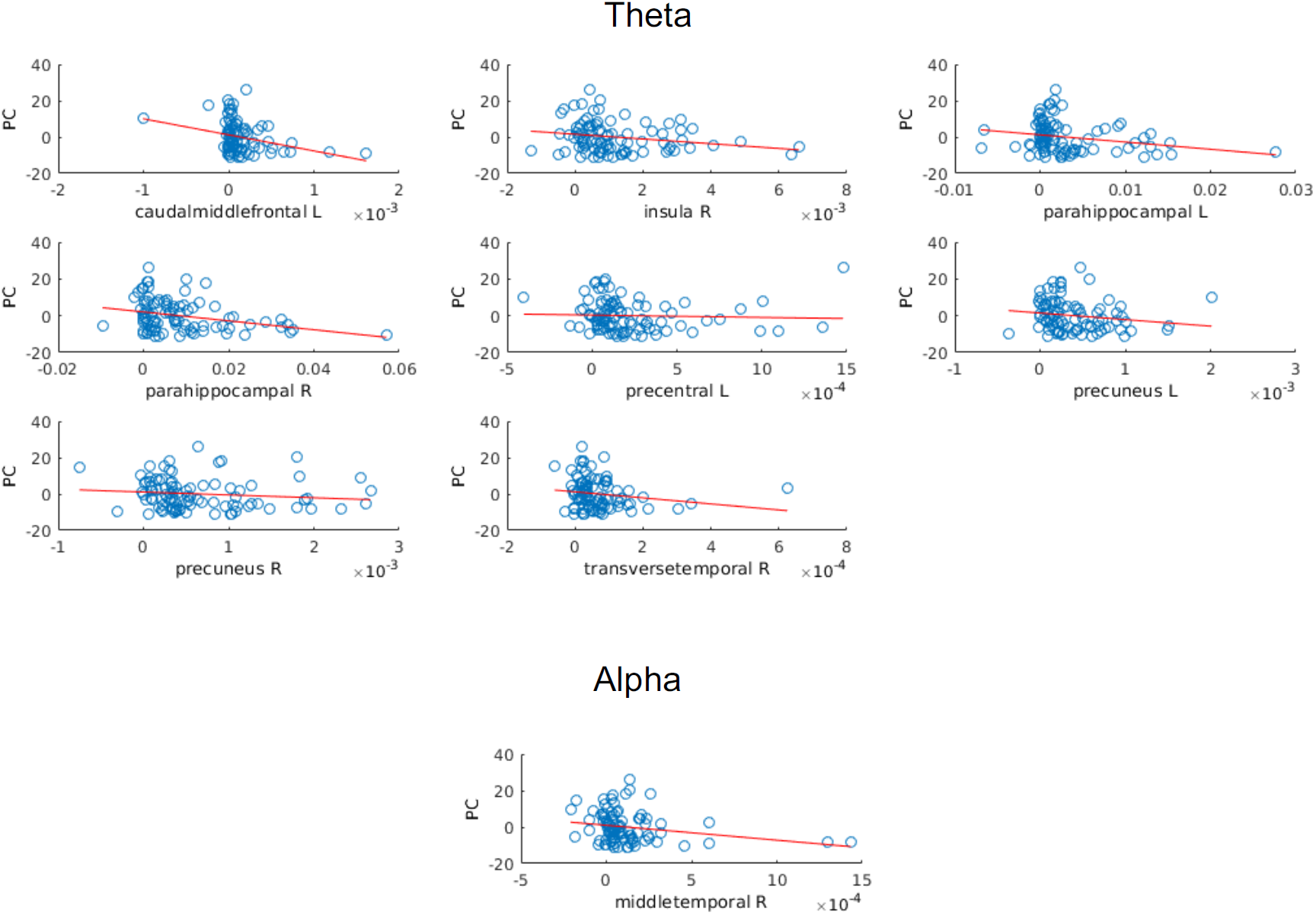
Neural symptom correlations. Significant correlations between the theta and alpha band global cognitive task-average activations and the top principal mental health symptom component (PC) are shown (Spearman correlations, p<0.05, fdr-corrected for multiple comparisons across ROIs and frequencies).

## References

Aftanas, L. I. and Golocheikine, S. A. (2001) ‘Human anterior and frontal midline theta and lower alpha reflect emotionally positive state and internalized attention: high-resolution EEG investigation of meditation’, Neuroscience Letters, 310(1), pp. 57–60. doi: 10.1016/S0304-3940(01)02094-8.

Anderson, T. W., Darling, D. A., Anderson, T. W. and Darling, D. A. (1952) ‘Asymptotic theory of certain goodness-of-fit criteria based on stochastic processes.’ Available at: https://www.scienceopen.com/document?vid=dd93f22b-2a06-4923-96bb-540c099e7db7 (Accessed: 10 April 2020).

Aron, A. R. (2007) ‘The neural basis of inhibition in cognitive control’, Neuroscientist, 13, pp. 214–228. doi: 10.1177/1073858407299288.

Aron, A. R. and Poldrack, R. A. (2005) ‘The Cognitive Neuroscience of Response Inhibition?: Relevance for Genetic Research in Attention-Deficit /’, Biological psychiatry, 57, pp. 1285–1292. doi: 10.1016/j.biopsych.2004.10.026.

Avissar, M., Powell, F., Ilieva, I., Respino, M., Gunning, F. M., Liston, C. and Dubin, M. J. (2017) ‘Functional Connectivity of the Left DLPFC to Striatum Predicts Treatment Response of Depression to TMS’, Brain stimulation, 10(5), pp. 919–925. doi: 10.1016/j.brs.2017.07.002.

Badre, D. (2011) ‘Defining an Ontology of Cognitive Control Requires Attention to Component Interactions’, Topics in cognitive science, 3, pp. 217–221. doi: 10.1111/j.1756-8765.2011.01141.x.

Barlow, H. B., Lal, S., Longuet-Higgins, H. C. and Sutherland, N. S. (1980) ‘The absolute efficiency of perceptual decisions’, Philosophical Transactions of the Royal Society of London. B, Biological Sciences. Royal Society, 290(1038), pp. 71–82. doi: 10.1098/rstb.1980.0083.

Bayerl, M., Dielentheis, T. F., Vucurevic, G., Gesierich, T., Vogel, F., Fehr, C., Stoeter, P., Huss, M. and Konrad, A. (2010) ‘Disturbed brain activation during a working memory task in drug-naive adult patients with ADHD’, NeuroReport, 21(6), pp. 442–446. doi: 10.1097/WNR.0b013e328338b9be.

Beltrán, D., Morera, Y., García-Marco, E. and Vega, M. de (2019) ‘Brain Inhibitory Mechanisms Are Involved in the Processing of Sentential Negation, Regardless of Its Content. Evidence From EEG Theta and Beta Rhythms’, Frontiers in Psychology, 10. doi: 10.3389/fpsyg.2019.01782.

Berryhill, M. E. and Olson, I. R. (2008) ‘The right parietal lobe is critical for visual working memory’, Neuropsychologia, 46(7), pp. 1767–1774. doi: 10.1016/j.neuropsychologia.2008.01.009.

Biederman, J., Makris, N., Valera, E. M., Monuteaux, M. C., Goldstein, J. M., Buka, S., Boriel, D. L., Bandyopadhyay, S., Kennedy, D. N., Caviness, V. S., Bush, G., Aleardi, M., Hammerness, P., Faraone, S. V. and Seidman, L. J. (2008) ‘Towards further understanding of the co-morbidity between attention deficit hyperactivity disorder and bipolar disorder: a MRI study of brain volumes’, Psychological Medicine, 38(7), pp. 1045–1056. doi: 10.1017/S0033291707001791.

Boudreau, B. and Poulin, C. (2008) ‘An examination of the validity of the Family Affluence Scale II (FAS II) in a general adolescent population of Canada’, Social Indicators Research, 94(1), p. 29. doi: 10.1007/s11205-008-9334-4.

Brass, M., Derrfuss, J., Forstmann, B. U. and von Cramon, D. Y. (2005) ‘The role of the inferior frontal junction area in cognitive control’, Trends in Cognitive Sciences, 9(7), pp. 314–316. doi: 10.1016/j.tics.2005.05.001.

Brzezicka, A. (2013) ‘Integrative deficits in depression and in negative mood states as a result of fronto-parietal network dysfunctions.’, Acta neurobiologiae experimentalis.

Bush, G. (2011) ‘Cingulate, Frontal, and Parietal Cortical Dysfunction in Attention-Deficit/Hyperactivity Disorder’, Biological Psychiatry. (Prefrontal Cortical Circuits Regulating Attention, Behavior and Emotion), 69(12), pp. 1160–1167. doi: 10.1016/j.biopsych.2011.01.022.

Buzsáki, G. and Moser, E. I. (2013) ‘Memory, navigation and theta rhythm in the hippocampal-entorhinal system’, Nature Neuroscience, 16(2), pp. 130–138. doi: 10.1038/nn.3304.

Canbeyli, R. (2010) ‘Sensorimotor modulation of mood and depression: An integrative review’, Behavioural Brain Research, 207(2), pp. 249–264. doi: 10.1016/j.bbr.2009.11.002.

Chambers, C. D., Garavan, H. and Bellgrove, M. A. (2009) ‘Insights into the neural basis of response inhibition from cognitive and clinical neuroscience’, Neuroscience and Biobehavioral Reviews, 33, pp. 631–646. doi: 10.1016/j.neubiorev.2008.08.016.

Chong, T. T. J., Williams, M. A., Cunnington, R., & Mattingley, J. B. (2008). Selective attention modulates inferior frontal gyrus activity during action observation. Neuroimage, 40(1), 298–307.

Christie, G. J. and Tata, M. S. (2009) ‘Right frontal cortex generates reward-related theta-band oscillatory activity’, Neuroimage, 48(2), pp. 415–422.

Cohen, J., Cohen, R., Adad, J., Cohen, J. M., Cohen, J. A., Mansfield, J. and Cohen, J. B. (1988) ‘Statistical power analysis for the behaviorla sciences’. Available at: https://www.scienceopen.com/document?vid=94bc2633-7c1a-41a3-89ee-56e75d596213 (Accessed: 10 April 2020).

Deckersbach, T., Dougherty, D. D. and Rauch, S. L. (2006) ‘Functional Imaging of Mood and Anxiety Disorders’, Journal of Neuroimaging, 16(1), pp. 1–10. doi: 10.1177/1051228405001474.

Delorme, A. and Makeig, S. (2004) ‘EEGLAB: an open source toolbox for analysis of single-trial EEG dynamics including independent component analysis’, Journal of Neuroscience Methods, 134(1), pp. 9–21. doi: 10.1016/j.jneumeth.2003.10.009.

Desikan, R. S., Ségonne, F., Fischl, B., Quinn, B. T., Dickerson, B. C., Blacker, D., Buckner, R. L., Dale, A. M., Maguire, R. P., Hyman, B. T., Albert, M. S. and Killiany, R. J. (2006) ‘An automated labeling system for subdividing the human cerebral cortex on MRI scans into gyral based regions of interest’, NeuroImage, 31(3), pp. 968–980. doi: 10.1016/j.neuroimage.2006.01.021.

Dosenbach, N. U. F., Fair, D. A., Miezin, F. M., Cohen, A. L., Wenger, K. K., Dosenbach, R. A. T., Fox, M. D., Snyder, A. Z., Vincent, J. L., Raichle, M. E., Schlaggar, B. L. and Petersen, S. E. (2007) ‘Distinct brain networks for adaptive and stable task control in humans’, Proceedings of the National Academy of Sciences. National Academy of Sciences, 104(26), pp. 11073–11078. doi: 10.1073/pnas.0704320104.

Efron, B. (1982). “The Jackknife, the bootstrap, and other resampling plans,” in CBMS-NSF Regional Conference Series in Applied Mathematics, Monograph 38 (Philadelphia, PA: SIAM). doi: 10.1137/1.9781611970319

Ekstrom, A. D., Caplan, J. B., Ho, E., Shattuck, K., Fried, I. and Kahana, M. J. (2005) ‘Human hippocampal theta activity during virtual navigation’, Hippocampus, 15(7), pp. 881–889. doi: 10.1002/hipo.20109.

Eriksen, B. and Eriksen, C. W. (1974) ‘Effects of noise letters upon the identification of a target letter in a nonsearch task *’, Perception & Psychophysics, 16, pp. 143–149.

Etkin, A., Gyurak, A. and O’Hara, R. (2013) ‘A neurobiological approach to the cognitive deficits of psychiatric disorders’, Dialogues in Clinical Neuroscience, 15(4), pp. 419–429.

Gazzaley, A. and Nobre, A. C. (2012) ‘Top-down modulation: Bridging selective attention and working memory’, Trends in cognitive sciences, 16(2), pp. 129–135. doi: 10.1016/j.tics.2011.11.014.

Genovese, C. R., Lazar, N. A. and Nichols, T. (2002) ‘Thresholding of Statistical Maps in Functional Neuroimaging Using the False Discovery Rate’, NeuroImage, 15(4), pp. 870–878. doi: 10.1006/nimg.2001.1037.

George, M. S., Taylor, J. J. and Short, E. B. (2013) ‘The Expanding Evidence Base for rTMS Treatment of Depression’, Current opinion in psychiatry, 26(1), pp. 13–18. doi: 10.1097/YCO.0b013e32835ab46d.

Gray, J. R. (2004) ‘Integration of Emotion and Cognitive Control’, Current Directions in Psychological Science, 13, pp. 46–48.

Greenberg, L. M. and Waldman, I. D. (1993) ‘Developmental normative data on the test of variables of attention (T.O.V.A.)’, Journal of Child Psychology and Psychiatry, 34, pp. 1019–1030.

Grimm, S., Beck, J., Schuepbach, D., Hell, D., Boesiger, P., Bermpohl, F., Niehaus, L., Boeker, H. and Northoff, G. (2008) ‘Imbalance between Left and Right Dorsolateral Prefrontal Cortex in Major Depression Is Linked to Negative Emotional Judgment: An fMRI Study in Severe Major Depressive Disorder’, Biological Psychiatry, 63(4), pp. 369–376. doi: 10.1016/j.biopsych.2007.05.033.

Gudmundsson, S., Runarsson, T. P., Sigurdsson, S., Eiriksdottir, G. and Johnsen, K. (2007) ‘Reliability of quantitative EEG features’, Clinical Neurophysiology, 118(10), pp. 2162–2171. doi: 10.1016/j.clinph.2007.06.018.

Guse, B., Falkai, P. and Wobrock, T. (2010) ‘Cognitive effects of high-frequency repetitive transcranial magnetic stimulation: a systematic review’, Journal of Neural Transmission, 117(1), pp. 105–122. doi: 10.1007/s00702-009-0333-7.

Haldane, M., & Frangou, S. (2006). Functional neuroimaging studies in mood disorders. Acta Neuropsychiatrica, 18(2), 88–99.

Hampshire, A., Chamberlain, S. R., Monti, M. M., Duncan, J. and Owen, A. M. (2010) ‘The role of the right inferior frontal gyrus: inhibition and attentional control’, NeuroImage, 3(50), pp. 1313–1319. doi: 10.1016/j.neuroimage.2009.12.109.

Hedges, L. V. and Olkin, I. (1985) Statistical methods for meta-analysis. Academic Press. Available at: https://agris.fao.org/agris-search/search.do?recordID=US880855188 (Accessed: 9 May 2020).

Heeger, D. and Landy, M. (2009) ‘Signal detection theory’, Encyclopedia of perception. SAGE Publications, pp. 887–892.

Hesslinger, B., Tebartz van Elst, L., Thiel, T., Haegele, K., Hennig, J. and Ebert, D. (2002) ‘Frontoorbital volume reductions in adult patients with attention deficit hyperactivity disorder’, Neuroscience Letters, 328(3), pp. 319–321. doi: 10.1016/S0304-3940(02)00554-2.

Holmes, C. J., Hoge, R., Collins, L., Woods, R., Toga, A. W. and Evans, A. C. (1998) ‘Enhancement of MR Images Using Registration for Signal Averaging’, Journal of Computer Assisted Tomography, 22(2), pp. 324–333.

Hoy, K. E., Bailey, N., Michael, M., Fitzgibbon, B., Rogasch, N. C., Saeki, T. and Fitzgerald, P. B. (2016) ‘Enhancement of Working Memory and Task-Related Oscillatory Activity Following Intermittent Theta Burst Stimulation in Healthy Controls’, Cerebral Cortex. Oxford Academic, 26(12), pp. 4563–4573. doi: 10.1093/cercor/bhv193.

Insel, T., Cuthbert, B., Garvey, M., Heinssen, R., Pine, D. S., Quinn, K., Sanislow, C. and Wang, P. (2010) ‘Research Domain Criteria (RDoC): Toward a New Classification Framework for Research on Mental Disorders’, American Journal of Psychiatry. American Psychiatric Publishing, 167(7), pp. 748–751. doi: 10.1176/appi.ajp.2010.09091379.

Inzlicht, M., Bartholow, B. D. and Hirsh, J. B. (2015) ‘HHS Public Access’, Trends in cognitive sciences, 19, pp. 126–132. doi: 10.1016/j.tics.2015.01.004.Emotional.

Jung, T. P., Makeig, S., Humphries, C., Lee, T. W., Mckeown, M. J., Iragui, V., & Sejnowski, T. J. (2000). Removing electroencephalographic artifacts by blind source separation. Psychophysiology, 37(2), 163–178.

Khanna, P. and Carmena, J. M. (2017) ‘Beta band oscillations in motor cortex reflect neural population signals that delay movement onset’, eLife, 6. doi: 10.7554/eLife.24573.

Kothe, C., Medine, D., Boulay, C., Grivich, M., Stenner, T. (2019) ‘Lab Streaming Layer’ Copyright https://labstreaminglayer.readthedocs.io/

Kroenke, K., Spitzer, R. L. and Williams, J. B. W. (2001) ‘The PHQ-9: Validity of a Brief Depression Severity Measure’, Journal of general internal medicine, 16, pp. 606–613.

Lavie, N., Hirst, A. and Fockert, J. W. D. (2004) ‘Load Theory of Selective Attention and Cognitive Control’, Journal of Experimental Psychology: General, 133, pp. 339–354. doi: 10.1037/0096-3445.133.3.339.

Lenartowicz, A., Delorme, A., Walshaw, P. D., Cho, A. L., Bilder, R. M., McGough, J. J., McCracken, J. T., Makeig, S. and Loo, S. K. (2014) ‘Electroencephalography Correlates of Spatial Working Memory Deficits in Attention-Deficit/Hyperactivity Disorder: Vigilance, Encoding, and Maintenance’, Journal of Neuroscience, 34(4), pp. 1171–1182. doi: 10.1523/JNEUROSCI.1765-13.2014.

Lenartowicz, A., Kalar, D. J., Congdon, E. and Poldrack, A. (2010) ‘Towards an Ontology of Cognitive Control’, Topics in cognitive science, 2, pp. 678–692. doi: 10.1111/j.1756-8765.2010.01100.x.

López-martín, S., Albert, J., Fernández-jaén, A. and Carretié, L. (2013) ‘Emotional distraction in boys with ADHD: Neural and behavioral correlates’, Brain and cognition, 83, pp. 10–20. doi: 10.1016/j.bandc.2013.06.004.

López-Martín, S., Albert, J., Fernández-Jaén, A. and Carretié, L. (2015) ‘Emotional response inhibition in children with attention-deficit/hyperactivity disorder: neural and behavioural data’, Psychological Medicine, 45, pp. 2057–2071. doi: 10.1017/S0033291714003195.

Luna, B., Marek, S., Larsen, B., Tervo-Clemmens, B. and Chahal, R. (2015) ‘An integrative model of the maturation of cognitive control’, Annual review of neuroscience, 38, pp. 151–170. doi: 10.1126/science.1249098.Sleep.

Makeig, S., Westerfield, M., Jung, T.-P., Enghoff, S., Townsend, J., Courchesne, E. and Sejnowski, T. J. (2002) ‘Dynamic Brain Sources of Visual Evoked Responses’, Science. American Association for the Advancement of Science, 295(5555), pp. 690–694. doi: 10.1126/science.1066168.

Massat, I., Slama, H., Kavec, M., Linotte, S., Mary, A., Baleriaux, D., Metens, T., Mendlewicz, J. and Peigneux, P. (2012) ‘Working Memory-Related Functional Brain Patterns in Never Medicated Children with ADHD’, PLoS ONE, 7(11). doi: 10.1371/journal.pone.0049392.

McEvoy, L. (2000) ‘Test-retest reliability of cognitive EEG’, Clin Neurophysiol, 111, pp. 457–463. doi: 10.1016/S1388-2457(99)00258-8.

McNaughton, N. (1997) ‘Cognitive Dysfunction Resulting from Hippocampal Hyperactivity—A Possible Cause of Anxiety Disorder?’, Pharmacology Biochemistry and Behavior, 56(4), pp. 603–611. doi: 10.1016/S0091-3057(96)00419-4.

Menon, V. and Uddin, L. Q. (2010) ‘Saliency, switching, attention and control: a network model of insula function’, Brain structure & function, 214(5–6), pp. 655–667. doi: 10.1007/s00429-010-0262-0.

Mensen, A., Gorban, C., Niklaus, M., Kuske, E. and Khatami, R. (2014) ‘The effects of theta-burst stimulation on sleep and vigilance in humans’, Frontiers in Human Neuroscience. Frontiers, 8. doi: 10.3389/fnhum.2014.00420.

Millan, M. J., Agid, Y., Brüne, M., Bullmore, E. T., Carter, C. S., Clayton, N. S., Connor, R., Davis, S., Deakin, B., DeRubeis, R. J., Dubois, B., Geyer, M. A., Goodwin, G. M., Gorwood, P., Jay, T. M., Joëls, M., Mansuy, I. M., Meyer-Lindenberg, A., Murphy, D., Rolls, E., Saletu, B., Spedding, M., Sweeney, J., Whittington, M. and Young, L. J. (2012) ‘Cognitive dysfunction in psychiatric disorders: characteristics, causes and the quest for improved therapy’, Nature Reviews Drug Discovery. Nature Publishing Group, 11(2), pp. 141–168. doi: 10.1038/nrd3628.

Misra, A., Ojeda, A., Mishra, J. (2018) ‘BrainE: a digital platform for evaluating, engaging and enhancing brain function’, Regents of the University of California Copyright SD2018–816.

Mishra, J. (2019) ‘Translating Science to our global communities’, Elephant in the Lab doi.org/10.5281/zenodo.2619955. 2019

Mishra, J., Anguera, J. A., Ziegler, D. A. and Gazzaley, A. (2013) ‘A Cognitive Framework for Understanding and Improving Interference Resolution in the Brain’, Progress in brain research, 207, pp. 351–377. doi: 10.1016/B978-0-444-63327-9.00013-8.

Mishra, J., Martinez, A., Schroeder, C. E. and Hillyard, S. A. (2012) ‘Spatial Attention Boosts Short-Latency Neural Responses in Human Visual Cortex’, NeuroImage, 59(2), pp. 1968–1978. doi: 10.1016/j.neuroimage.2011.09.028.

Mishra, J., de Villers-Sidani, E., Merzenich, M. and Gazzaley, A. (2014) ‘Adaptive training diminishes distractibility in aging across species’, Neuron, 84(5), pp. 1091–1103.

Nee, D. E., Brown, J. W., Askren, M. K., Berman, M. G., Demiralp, E., Krawitz, A. and Jonides, J. (2013) ‘A Meta-analysis of Executive Components of Working Memory’, Cerebral Cortex, 23(2), pp. 264–282. doi: 10.1093/cercor/bhs007.

van Noordt, S. and Segalowitz, S. J. (2012) ‘Performance monitoring and the medial prefrontal cortex: a review of individual differences and context effects as a window on self-regulation’, Frontiers in Human Neuroscience. Frontiers, 6. doi: 10.3389/fnhum.2012.00197.

Nunez, P. L. (2010). REST: a good idea but not the gold standard. Clinical neurophysiology: official journal of the International Federation of Clinical Neurophysiology, 121(12), 2177.

Oberman, L., Edwards, D., Eldaief, M. and Pascual-Leone, A. (2011) ‘Safety of theta burst transcranial magnetic stimulation: a systematic review of the literature’, Journal of Clinical Neurophysiology, 28(1), p. 67.

Ojeda, A., Klug, M., Kreutz-Delgado, K., Gramann, K. and Mishra, J. (2019) ‘A Bayesian framework for unifying data cleaning, source separation and imaging of electroencephalographic signals’, bioRxiv. Cold Spring Harbor Laboratory, p. 559450. doi: 10.1101/559450v3.

Ojeda, A., Kreutz-Delgado, K. and Mullen, T. (2018) ‘Fast and robust Block-Sparse Bayesian learning for EEG source imaging’, Neuroimage, 174, pp. 449–462.

Okon-Singer, H., Hendler, T., Pessoa, L. and Shackman, A. J. (2015) ‘The neurobiology of emotion– cognition interactions: fundamental questions and strategies for future research’, Frontiers in Human Neuroscience. Frontiers, 9. doi: 10.3389/fnhum.2015.00058.

Palva, S. and Palva, J. M. (2007) ‘New vistas for α-frequency band oscillations’, Trends in Neurosciences, 30(4), pp. 150–158. doi: 10.1016/j.tins.2007.02.001.

Pascual-Marqui, R. D., Michel, C. M. and Lehmann, D. (1994) ‘Low resolution electromagnetic tomography: A new method for localizing electrical activity in the brain’, International Journal of Psychophysiology. Netherlands: Elsevier Science, 18(1), pp. 49–65. doi: 10.1016/0167-8760(84)90014-X.

Pessoa, L. (2009) ‘How do emotion and motivation direct executive control?’, Cell, 13, pp. 160–166. doi: 10.1016/j.tics.2009.01.006.

Pfurtscheller, G. (1999) ‘EEG event - related desynchronization (ERD) and event - releated synchronization (ERS).’, Electroencephalography: Basic Principles, Clinical Applications and Releated Fields. Clinical Applications and Related Fields 4th Edition, pp. 958–967.

Picazio, S., Veniero, D., Ponzo, V., Caltagirone, C., Gross, J., Thut, G. and Koch, G. (2014) ‘Prefrontal Control over Motor Cortex Cycles at Beta Frequency during Movement Inhibition’, Current Biology, 24(24), pp. 2940–2945. doi: 10.1016/j.cub.2014.10.043.

Posner, M. I. and Rothbart, M. K. (2009) ‘Toward A Physical Basis of Attention and Self Regulation’, Physics of life reviews, 6(2), pp. 103–120. doi: 10.1016/j.plrev.2009.02.001.

Price, J. L., & Drevets, W. C. (2012). Neural circuits underlying the pathophysiology of mood disorders. Trends in cognitive sciences, 16(1), 61–71.

Reinholdt-Dunne, M. L., Mogg, K. and Bradley, B. P. (2013) ‘Attention control: Relationships between self-report and behavioural measures, and symptoms of anxiety and depression’, Cognition and Emotion. Routledge, 27(3), pp. 430–440. doi: 10.1080/02699931.2012.715081.

Rossi, S., Hallett, M., Rossini, P. M. and Pascual-Leone, A. (2009) ‘Safety, ethical considerations, and application guidelines for the use of transcranial magnetic stimulation in clinical practice and research’, Clinical Neurophysiology, 120(12), pp. 2008–2039. doi: 10.1016/j.clinph.2009.08.016.

Rossi, S., Hallett, M., Rossini, P. M., Pascual-Leone, A. and Safety of TMS Consensus Group (2009) ‘Safety, ethical considerations, and application guidelines for the use of transcranial magnetic stimulation in clinical practice and research’, Clinical Neurophysiology, 120(12), pp. 2008–2039.

Sadaghiani, S., Dombert, P. L., Løvstad, M., Funderud, I., Meling, T. R., Endestad, T., Knight, R. T., Solbakk, A.-K. and D’Esposito, M. (2019) ‘Lesions to the Fronto-Parietal Network Impact Alpha-Band Phase Synchrony and Cognitive Control’, Cerebral Cortex. Oxford Academic, 29(10), pp. 4143–4153. doi: 10.1093/cercor/bhy296.

Sadaghiani, S., Scheeringa, R., Lehongre, K., Morillon, B., Giraud, A.-L., D’Esposito, M. and Kleinschmidt, A. (2012) ‘Alpha-Band Phase Synchrony Is Related to Activity in the Fronto-Parietal Adaptive Control Network’, Journal of Neuroscience. Society for Neuroscience, 32(41), pp. 14305–14310. doi: 10.1523/JNEUROSCI.1358-12.2012.

Shipstead, Z., Harrison, T. L. and Engle, R. W. (2012) ‘Working memory capacity and visual attention: Top-down and bottom-up guidance’, Quarterly Journal of Experimental Psychology, 65, pp. 401–407. doi: 10.1080/17470218.2012.655698.

Siegle, G. J., Thompson, W., Carter, C. S., Steinhauer, S. R. and Thase, M. E. (2007) ‘Increased Amygdala and Decreased Dorsolateral Prefrontal BOLD Responses in Unipolar Depression: Related and Independent Features’, Biological Psychiatry, 61(2), pp. 198–209. doi: 10.1016/j.biopsych.2006.05.048.

Song, S., Zilverstand, A., Song, H., Uquillas, F. d’Oleire, Wang, Y., Xie, C., Cheng, L. and Zou, Z. (2017) ‘The influence of emotional interference on cognitive control: A meta-analysis of neuroimaging studies using the emotional Stroop task’, Scientific Reports. Nature Publishing Group, 7(1), pp. 1–9. doi: 10.1038/s41598-017-02266-2.

Spearman, C. (1904) ‘“The proof and measurement of association between two things”‘. Available at:https://www.scienceopen.com/document?vid=1b6909e5-63d2-43b0-a160-84f677829b20 (Accessed: 10 April 2020).

Spitzer, R. L., Kroenke, K., Williams, J. B. W. and Loewe, B. (2006) ‘A Brief Measure for Assessing Generalized Anxiety Disorder: The GAD-7’, Archives of internal medicine, 166, pp. 1092–1097.

Sternberg, S. (1966) ‘High-speed scanning in human memory’, science, 153, pp. 652–654.

Sylvester, C. M., Corbetta, M., Raichle, M. E., Rodebaugh, T. L., Schlaggar, B. L., Sheline, Y. I., Zorumski, C. F. and Lenze, E. J. (2012) ‘Functional network dysfunction in anxiety and anxiety disorders’, Trends in Neurosciences, 35(9), pp. 527–535. doi: 10.1016/j.tins.2012.04.012.

Tettamanti, M., Manenti, R., Della Rosa, P. A., Falini, A., Perani, D., Cappa, S. F. and Moro, A. (2008) ‘Negation in the brain: Modulating action representations’, NeuroImage, 43(2), pp. 358–367. doi: 10.1016/j.neuroimage.2008.08.004.

Thai, N., Taber-Thomas, B. C. and Pérez-Edgar, K. E. (2016) ‘Neural correlates of attention biases, behavioral inhibition, and social anxiety in children: An ERP study’, Developmental Cognitive Neuroscience, 19, pp. 200–210. doi: 10.1016/j.dcn.2016.03.008.

Thimm, A. and Funke, K. (2015) ‘Multiple blocks of intermittent and continuous theta-burst stimulation applied via transcranial magnetic stimulation differently affect sensory responses in rat barrel cortex’, The Journal of Physiology, 593(4), pp. 967–985. doi: 10.1113/jphysiol.2014.282467.

Tottenham, N., Tanaka, J. W., Leon, A. C., McCarry, T., Nurse, M., Hare, T. A., Marcus, D. J., Westerlund, A., Casey, B. J. and Nelson, C. (2009) ‘The NimStim set of facial expressions: Judgments from untrained research participants’, Psychiatry Research, 168, pp. 242–249. doi: 10.1016/j.psychres.2008.05.006.

The. Vandierendonck, A. (2017) ‘A comparison of methods to combine speed and accuracy measures of performance: A rejoinder on the binning procedure’, Behavior Research Methods, 49(2), pp. 653–673. doi: 10.3758/s13428-016-0721-5.

Vékony, T., Németh, V. L., Holczer, A., Kocsis, K., Kincses, Z. T., Vécsei, L. and Must, A. (2018) ‘Continuous theta-burst stimulation over the dorsolateral prefrontal cortex inhibits improvement on a working memory task’, Scientific Reports, 8. doi: 10.1038/s41598-018-33187-3.

Verbruggen, F., Aron, A. R., Stevens, M. A. and Chambers, C. D. (2010) ‘Theta burst stimulation dissociates attention and action updating in human inferior frontal cortex’, Proceedings of the National Academy of Sciences. National Academy of Sciences, 107(31), pp. 13966–13971. doi: 10.1073/pnas.1001957107.

Viejo-Sobera, R., Redolar-Ripoll, D., Boixadós, M., Palaus, M., Valero-Cabré, A. and Marron, E. M. (2017) ‘Impact of Prefrontal Theta Burst Stimulation on Clinical Neuropsychological Tasks’, Frontiers in Neuroscience, 11. doi: 10.3389/fnins.2017.00462.

W, P. (2005) ‘Toward establishing a therapeutic window for rTMS by theta burst stimulation.’, Neuron, 45(2), pp. 181–183. doi: 10.1016/j.neuron.2005.01.008.

Waugh, C. E., Lemus, M. G. and Gotlib, I. H. (2014) ‘The role of the medial frontal cortex in the maintenance of emotional states’, Social Cognitive and Affective Neuroscience, 9(12), pp. 2001–2009. doi: 10.1093/scan/nsu011.

Wodka, E. L., Mahone, E. M., Blankner, J. G., Larson, J. C. G., Fotedar, S., Denckla, M. B. and Mostofsky, S. H. (2007) ‘Evidence that response inhibition is a primary deficit in ADHD’, Journal of Clinical and Experimental Neuropsychology. Routledge, 29(4), pp. 345–356. doi: 10.1080/13803390600678046.

Zanto, T. P., Rubens, M. T., Thangavel, A. and Gazzaley, A. (2011) ‘Causal role of the prefrontal cortex in top-down modulation of visual processing and working memory’, Nature neuroscience, 14(5), pp. 656–661. doi: 10.1038/nn.2773.

Zavala, B., Jang, A., Trotta, M., Lungu, Codrin I., Brown, P. and Zaghloul, K. A. (2018) ‘Cognitive control involves theta power within trials and beta power across trials in the prefrontal-subthalamic network’, Brain: A Journal of Neurology, 141(12), pp. 3361–3376. doi: 10.1093/brain/awy266.

Zavala, B., Jang, A., Trotta, M., Lungu, Codrin I, Brown, P. and Zaghloul, K. A. (2018) ‘Cognitive control involves theta power within trials and beta power across trials in the prefrontal-subthalamic network’, Brain, 141(12), pp. 3361–3376. doi: 10.1093/brain/awy266.

Zhao, X.-H., Wang, P.-J., Li, C.-B., Hu, Z.-H., Xi, Q., Wu, W.-Y. and Tang, X.-W. (2007) ‘Altered default mode network activity in patient with anxiety disorders: An fMRI study’, European Journal of Radiology. (Prostate), 63(3), pp. 373–378. doi: 10.1016/j.ejrad.2007.02.006.

